# NeuroPupil: A generalization-first framework for scalable and biologically informative cross-species pupillometry

**DOI:** 10.64898/2026.05.14.725098

**Authors:** Kemal Ozdemirli, Tenesha Connor, Berfin Dinc, Kaleb Kim, Macit Emre Lacin, Miguel Maldonado, Shreya Guha, Kiersten Hawk, Caglar Oksuz, Frederick Bell, Zhiyao Zhang, Thiago Peixoto Leal, Nicholas Sarn, Anthony Sloan, Anthony Chomyk, Fatema Ghasia, Bruce Trapp, Ignacio Mata, Justin D. Lathia, Charis Eng, Murat Yildirim

## Abstract

Quantitative pupillometry provides a noninvasive window into brain state and neurological function, but its broader use across experimental and clinical settings is limited by challenges in achieving accurate, scalable, and generalizable measurements. Here, we present NeuroPupil, a deep learning framework for high-throughput, cross-species pupillometry that emphasizes robust generalization across subjects, behavioral contexts, and imaging conditions. Through systematic benchmarking of training strategies and network architectures, we identify pooled multi-subject training combined with an optimized U-Net architecture as a key determinant of reliable and transferable pupil tracking performance.

Across diverse mouse and human datasets, NeuroPupil achieves improved accuracy and substantial gains in computational efficiency compared to existing approaches, enabling practical analysis of large-scale datasets. We further demonstrate that improved pupil tracking fidelity enhances downstream biological inference: NeuroPupil-derived pupil features significantly improve prediction of distributed cortical activity in behaving mice and preserve diagnostically relevant temporal structure in human clinical recordings. These findings highlight the importance of precise and scalable measurement for linking pupil dynamics to brain activity and clinical phenotypes.

By integrating benchmarking, scalability, and accessible software tools, NeuroPupil provides a reproducible framework for large-scale pupillometry and facilitates its application in systems and translational neuroscience.

## INTRODUCTION

Pupillometry offers a uniquely accessible and noninvasive readout of internal brain state, capturing fluctuations in autonomic tone, arousal, and neuromodulatory activity across species^1–3^. As a result, pupil dynamics are increasingly used in both basic neuroscience and clinical research to probe cognitive processing, neurological disease, and brain–behavior relationships^4,5^. Despite this growing importance, widely used general-purpose pose-estimation frameworks applied to pupillometry, such as DeepLabCut^6^ and Facemap^7^, exhibit limitations in scalability, consistency, and generalization across experimental conditions.

In clinical contexts, subtle changes in pupil dynamics, including altered baseline diameter, constriction velocity, and temporal variability, have been associated with a wide range of neurological disorders, such as traumatic brain injury, Parkinson’s disease, autism spectrum disorder, and multiple sclerosis^8–14^. In preclinical neuroscience, pupil measurements are routinely combined with electrophysiology and large-scale calcium imaging to infer global brain state and neuromodulatory tone during behavior^15–17^. Together, these applications place stringent demands on pupillometry methods, requiring high temporal precision, robustness to motion and occlusion, and consistency across subjects, sessions, and recording environments.

Existing pupillometry approaches fall short of these requirements. Manual pupil examination lacks sensitivity and is prone to high inter-observer variability, particularly for small or rapid pupil changes^18^. Automated infrared pupillometers improve objectivity but are limited to specialized hardware and constrained experimental settings^19^. More recently, deep learning–based methods such as DeepLabCut and Facemap have enabled automated pupil tracking from video data in both humans and animals^7,20^. While transformative, these tools typically rely on per-session or per-subject model training, require substantial manual annotation, and scale poorly to large, heterogeneous datasets^7,20,21^.

Consequently, performance often degrades under variable imaging conditions, disease-related changes in pupil morphology, or deployment across laboratories and species.

A key challenge in computational pupillometry is identifying training strategies and model architectures that optimize generalization across animals, sessions, behaviors, and clinical contexts while maintaining high accuracy and computational efficiency. Although recent advances in pose estimation and behavioral tracking have begun to explore data aggregation and foundation-style modeling as routes toward improved generalization^22,23^, these principles have not been systematically evaluated or operationalized for pupillometry. This gap is particularly consequential because small, systematic errors in pupil boundary localization can propagate into substantial biases in downstream neural decoding, spectral analysis, and clinical classification.

Here, we present NeuroPupil, a deep learning framework built on the SLEAP pose estimation platform that addresses these limitations by explicitly prioritizing generalization and scalability. Through systematic comparison of training strategies, network architectures, and annotation schemes, we identify the combination of pooled multi-subject training and an optimized U-Net–based architecture as the dominant determinant of robust and generalizable pupil tracking, with substantially greater impact than annotation density alone. Building on this insight, we develop a unified pupillometry pipeline that integrates optimized training and inference strategies with user-friendly graphical user interfaces and supports flexible deployment on CPU, GPU, and high-performance computing architectures. This design enables high-precision tracking with sparse annotations, robust generalization across preclinical disease models and heterogeneous human clinical datasets, and efficient scaling from desktop environments to large-scale computational infrastructure.

Using NeuroPupil, we further demonstrate that improvements in pupil tracking fidelity directly enhance downstream biological inference. NeuroPupil-derived pupil features enable more accurate prediction of distributed cortical activity in behaving mice and preserve fine-scale temporal structure that supports robust analysis of disease-associated pupil dynamics in human clinical data. By reframing pupillometry as a generalization-first measurement problem and embedding this principle into an end-to-end, reproducible workflow, NeuroPupil advances pupillometry from an ad hoc analytical step to a scalable, biologically informative measurement suitable for integrative and translational neuroscience.

## RESULTS

### NeuroPupil: A Systematic Benchmarking–Driven Framework for Generalizable, High-Precision Pupillometry Across Behavioral and Disease Contexts

Accurate and scalable pupillometry is essential for quantifying arousal, neuromodulatory tone, and cognitive state in behaving animals, yet remains technically challenging due to variability in imaging resolution, behavioral dynamics, inter-animal differences, and disease-associated alterations in pupil morphology. Although deep learning–based tracking tools are widely used, existing workflows typically rely on per-session or per-animal models, lack principled guidance for training strategy selection, and scale poorly to large datasets. To address these limitations, we developed NeuroPupil and used it as a testbed to systematically evaluate training strategies and model architectures for scalable pupillometry.

We began by performing a comprehensive head-to-head benchmarking of pupillometry models across two widely used behavioral paradigms that differ substantially in imaging resolution and motion statistics: a low-resolution wheel-running setup (**Fig. 1A, left**) and a high-resolution airball setup (**Fig. 1A, right**). In both paradigms, pupil dynamics were recorded using infrared illumination to generate high-contrast pupil boundaries during ongoing behavior. This design enabled direct evaluation of model generalization across distinct optical and behavioral regimes.

**Figure 1.**
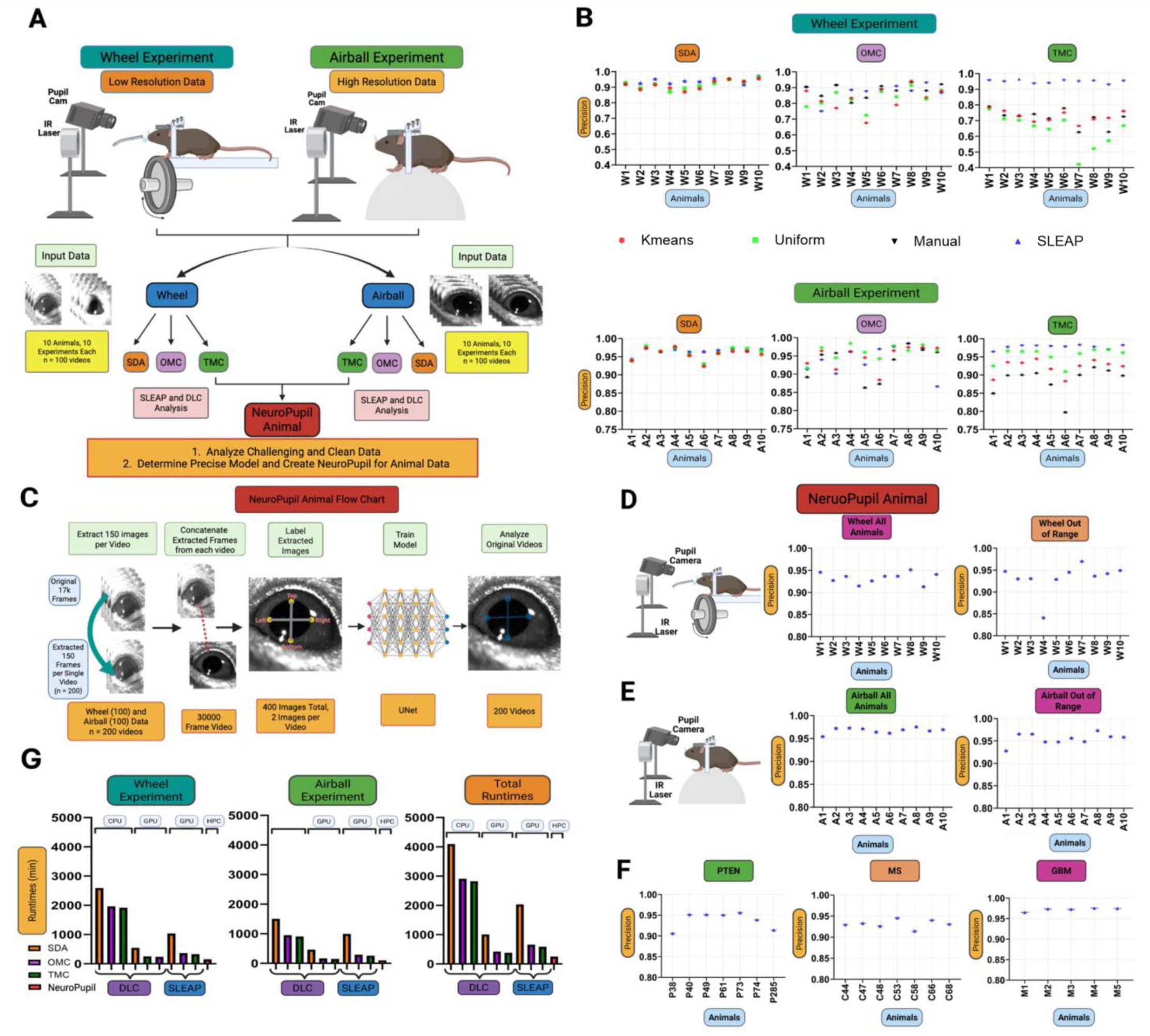
NeuroPupil enables high-precision, generalizable, and ultrafast pupillometry across mouse experiments. (**A**) Experimental paradigms and modeling strategies for mouse pupillometry acquired under infrared illumination from a low-resolution wheel-running setup (left) and a high-resolution airball setup (right). Pupil data were analyzed using three training strategies: Single-Day Animal (SDA), in which one model is trained per video; One-Mouse Concatenation (OMC), in which all sessions from a single mouse are pooled into one model; and Ten-Mouse Concatenation (TMC), in which data from multiple mice are combined to train a single model per experimental condition. Each strategy was implemented using DeepLabCut (ResNet-50 backbone) and SLEAP (U-Net backbone). Benchmarking identified SLEAP-TMC as the most precise and robust configuration across experimental contexts, motivating the development of a global pupillometry model, termed NeuroPupil. (**B**) Precision benchmarking across training strategies and frameworks. Precision values are shown for SDA, OMC, and TMC models trained with DeepLabCut and SLEAP for both wheel (top) and airball (bottom) datasets. Across animals and experimental paradigms, TMC-trained models consistently outperformed SDA and OMC approaches, with SLEAP-TMC achieving the highest and most stable precision. (**C**) NeuroPupil training and analysis workflow. From each video (n = 200; 100 wheel and 100 airball), 150 frames were uniformly sampled and concatenated using a custom graphical user interface. A total of 400 images (two per video) were manually annotated with four pupil boundary keypoints (top, bottom, left, right) and used to train a U-Net–based SLEAP model. The trained NeuroPupil model was subsequently applied to full video datasets for automated, frame-by-frame pupil tracking at scale. (**D**) NeuroPupil precision across all animals in wheel-running experiments. Performance is shown for all animals included in training and for “out-of-range” cases characterized by extreme pupil dilation or constriction and challenging imaging conditions, demonstrating robust performance across pupil states. (E) NeuroPupil precision across all animals in airball experiments. As in wheel recordings, NeuroPupil maitains high precision across animals and generalizes effectively to out-of-range pupil dynamics in high-resolution imaging conditions. (F) Generalization of NeuroPupil across disease models. Precision is shown for cohorts modeling PTEN-associated autism spectrum disorder, cuprizone-induced multiple sclerosis (MS), and glioblastoma (SB28 GBM), demonstrating robust performance despite disease-related changes in pupil dynamics and facial morphology. (**G**) Runtime benchmarking of pupillometry pipelines. Processing times are compared across DeepLabCut (SDA, OMC, TMC), SLEAP (SDA, OMC, TMC), and NeuroPupil deployed on high-performance computing (HPC) infrastructure. NeuroPupil achieves the fastest runtimes for both wheel and airball datasets, enabling efficient analysis of large-scale pupillometry data without sacrificing precision.

A central conceptual component of NeuroPupil is the explicit evaluation of data aggregation as a design principle for pupillometry. NeuroPupil is implemented on the SLEAP pose-estimation framework and uses this platform as a unified testbed to assess how different training strategies affect robustness and generalization. We compared three training strategies representing increasing levels of generalization: Single-Day Animal (SDA) training, in which each video is modeled independently; One-Mouse Concatenation (OMC), in which all sessions from a single animal are pooled together into a model; and Ten-Mouse Concatenation (TMC), in which data from multiple animals under the same experimental condition are combined into a single model. To disentangle the effects of training strategy from model architecture, we implemented each strategy using both SLEAP (U-Net backbone) and DeepLabCut (ResNet-50 backbone) as parallel benchmarking frameworks (**Supp. Fig. 1**).

To ensure that observed performance differences reflected methodological choices rather than annotation bias, we first optimized and standardized frame-selection procedures. Using the wheel dataset, we compared manual frame selection with automated approaches, including k-means clustering and uniform sampling, across a range of labeled frame counts using DeepLabCut (**Supp. Fig. 2**). Precision was comparable across selection methods, and increasing manual labeling density yielded minimal performance gains. These results indicate that sparse, automatically selected annotations are sufficient for accurate pupil tracking, motivating their use in subsequent NeuroPupil training and benchmarking experiments.

We next performed systematic optimization of SLEAP parameters for pupillometry analysis, including clustering configuration, tracking algorithm, instance-matching strategy, and assignment method (**Supp. Fig. 3**). Optimal performance was achieved using compact, information-rich training sets combined with flow-based tracking, centroid matching, and greedy assignment. Importantly, this analysis revealed that training strategy and data diversity, not annotation volume or network complexity, are the dominant determinants of pupillometry performance.

Across both imaging paradigms, TMC-trained models consistently outperformed SDA (**Supp. Fig. 4**) and OMC (**Supp. Fig. 5**) approaches, achieving higher precision and reduced inter-animal variability in both wheel (top) and airball (bottom) experiments (**Fig. 1B; Supp. Fig. 6**). This effect was most pronounced for SLEAP-based models, indicating that pooled multi-animal training enables the network to learn invariant representations of pupil geometry that generalize across behavioral states, imaging resolutions, and animals. Using two representative wheel-running datasets, one challenging (W9) and one clean (W3), we further showed that labeling as few as two frames per video was sufficient to achieve optimal performance (**Supp. Fig. 7**). For image-labeling comparisons with SLEAP, we used only the DeepLabCut Manual Strategy, labeling the identical frames selected for the SLEAP model to ensure direct methodological equivalence. We did not include DeepLabCut k-means or uniform frame selection for this comparison (described more detail in the DeepLabCut OMC and TMC Methods Section), as these approaches do not enforce equal sampling across videos and therefore do not provide a one-to-one correspondence (e.g., two frames per video). Since the SLEAP training procedure selects an equal number of frames per video, DeepLabCut Manual labeling was the only strategy that enabled a controlled and unbiased comparison. Collectively, these results establish pooled multi-animal training as the core design principle underlying NeuroPupil.

Guided by these benchmarking results, we instantiated NeuroPupil as a global pupillometry model trained using the TMC strategy on pooled low- and high-resolution datasets (**Fig. 1C**). NeuroPupil was trained on a deliberately sparse yet diverse dataset comprising 400 annotated images drawn from 200 videos spanning two behavioral paradigms, demonstrating that high-performance pupillometry can be achieved with minimal manual annotation when training data are structured for generalization. Once trained, NeuroPupil enables fully automated, frame-by-frame pupil tracking across large video collections.

NeuroPupil maintained high precision across all animals and experimental conditions, including challenging “out-of-range” wheel and airball cases characterized by extreme pupil dilation or constriction, motion artifacts, and partial occlusions (**Fig. 1D–E**; **Supp. Figs. 8–9**). Notably, performance generalized across multiple disease models, including PTEN-associated autism spectrum disorder (**Supp. Fig. 10**), cuprizone-induced multiple sclerosis (**Supp. Fig. 11**), and glioblastoma following SB28 tumor injection (**Supp. Fig. 12**), underscoring NeuroPupil’s robustness to disease-related changes in pupil dynamics (**Fig. 1F**). The “out-of-range” and disease animals were derived from test videos not used during the NeuroPupil mouse model training, demonstrating the robust out of sample generalization.

In addition to improved accuracy and robustness, NeuroPupil provides substantial computational advantages. Runtime analysis revealed that NeuroPupil markedly outperforms conventional DeepLabCut- and SLEAP-based workflows, particularly when deployed on high-performance computing infrastructure (HPC) (**Fig. 1G; Supp. Fig. 13**). To broaden accessibility, we developed a modular, open-source graphical user interface framework composed of two complementary tools, mediaGUI for video preprocessing and concatenation and sleapGUI Pupil for large-scale pupillometry model inference, analysis, and data export, which are designed to be used sequentially within a unified workflow (**Supp. Fig. 13**). The framework enables laboratories without access to high-performance computing resources to train custom NeuroPupil models, perform large-scale analyses beyond the capabilities of the default SLEAP GUI, and seamlessly extract SLEAP (.slp) files, CSVs of labeled x,y coordinates, and labeled videos (**Supp. Fig. 13**). This acceleration enables rapid analysis of datasets comprising millions of frames, transforming pupillometry from a technical bottleneck into a scalable, high-throughput measurement.

Together, these results introduce NeuroPupil as a principled, data-efficient, and computationally scalable pupillometry framework. By identifying pooled multi-animal training as the dominant factor governing robust pupil tracking and embedding this insight into an end-to-end workflow, NeuroPupil provides a generalizable solution that advances pupillometry from an ad hoc analysis step to a reproducible and field-ready measurement for basic and translational neuroscience.

### NeuroPupil achieves lower pupil estimation error and orders-of-magnitude speedup compared to Facemap

To benchmark NeuroPupil against an established pupillometry approach, we directly compared its pupil size estimates and computational performance with those obtained using Facemap across both low-resolution wheel-running (**Supp. Videos 1-2**) and high-resolution airball datasets (**Supp. Videos 3-4**) (**Fig. 2**). Representative time series illustrate that NeuroPupil closely tracks pupil dynamics while exhibiting reduced frame-to-frame noise and fewer spurious fluctuations relative to Facemap across behavioral contexts and imaging resolutions (**Fig. 2A–D**).

**Figure 2.**
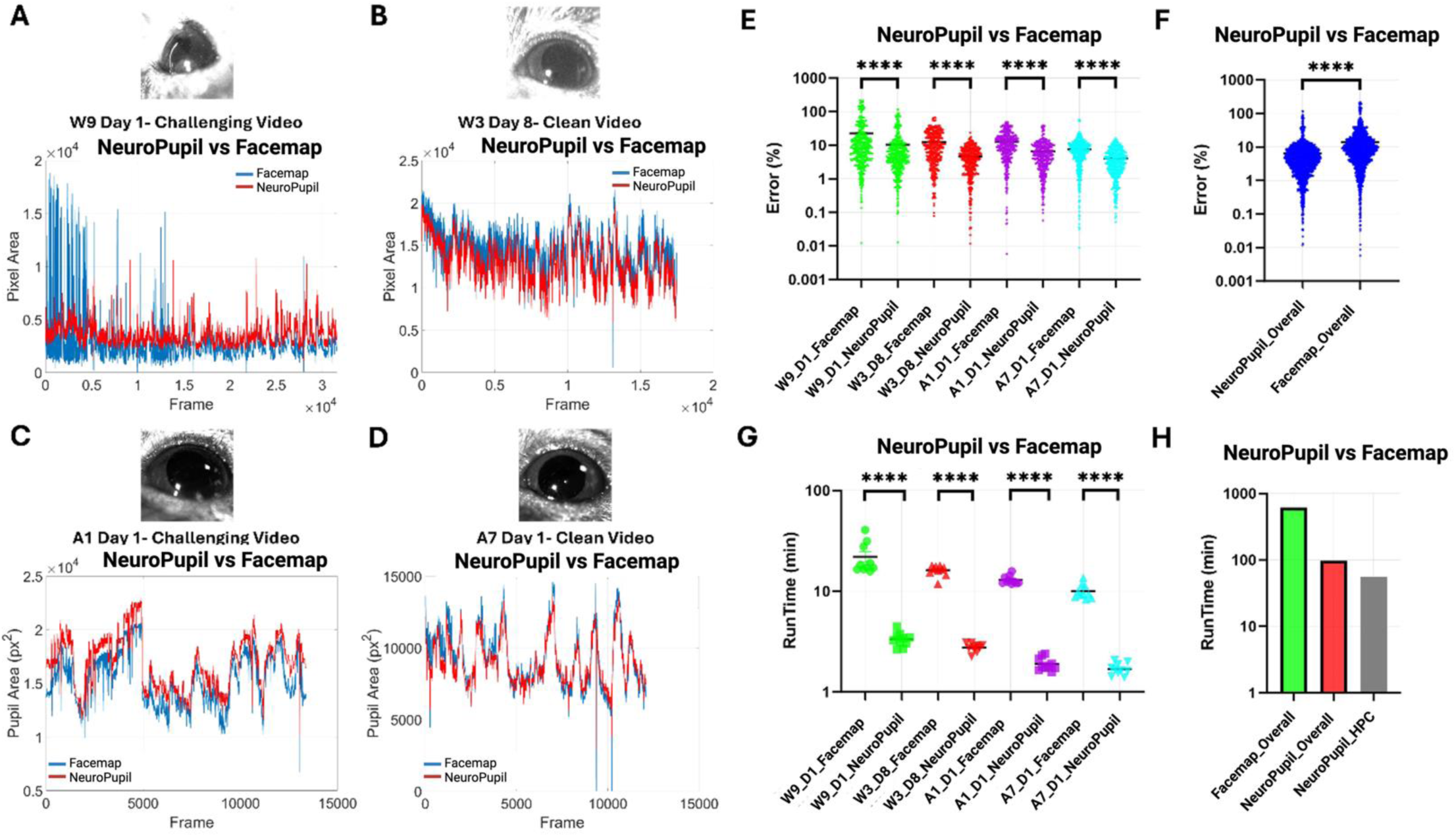
NeuroPupil outperforms Facemap in pupil size accuracy and computational efficiency across behavioral paradigms. (A–D) Representative pupil area time series comparing Facemap (blue) and NeuroPupil (red) across low-resolution wheel-running and high-resolution airball experiments, shown for individual animals and sessions: **(A)** W9, day 1; **(B)** W3, day 8; **(C)** A1, day 1; and **(D)** A7, day 1. Insets show representative eye images for each dataset. **(E) Percent error of pupil area estimates relative to human-annotated ground truth.** Ten frames per video were manually annotated by four independent expert annotators. Error distributions are shown separately for each animal (W9, W3, A1, A7), comparing Facemap and NeuroPupil. NeuroPupil exhibits significantly lower error across all animals (two-sided t-test, ****p < 0.0001). **(F) Overall percent error aggregated across all animals and sessions.** NeuroPupil achieves significantly lower error relative to Facemap when compared against human annotations (two-sided t-test, ****p < 0.0001). **(G) Runtime comparisons for FaceMap and NeuroPupil for individual videos across animals (W9, W3, A1, A7)**. NeuroPupil consistently achieves substantially faster processing times across datasets (two-sided t-test, ****p < 0.0001). **(H) Overall runtime comparison across all videos and animals**. Processing times are shown for Facemap executed on a local workstation, NeuroPupil executed on a local workstation, and NeuroPupil deployed on high-performance computing (HPC) infrastructure. NeuroPupil demonstrates marked reductions in runtime, with further acceleration when deployed on HPC.

To quantitatively assess accuracy, we generated human-annotated ground truth by manually labeling pupil boundaries in ten randomly selected frames per video using four independent expert annotators. Percent error was computed relative to consensus annotations. Across all tested animals and sessions, NeuroPupil consistently achieved significantly lower error than Facemap (**Fig. 2E**), with improvements observed in both wheel (W9, W3) and airball (A1, A7) recordings. When aggregated across animals, NeuroPupil exhibited a robust reduction in overall percent error relative to Facemap (**Fig. 2F**; two-sided t-test, ****p < 0.0001), indicating improved agreement with expert annotations across imaging conditions.

Beyond accuracy, we evaluated computational efficiency. Runtime analyses on the local computer showed that NeuroPupil processed individual videos substantially faster than Facemap across all animals and experimental conditions (**Fig. 2G**). Importantly, because Facemap relies on a pretrained model, video specific configuration that requires tedious manual adjustment of pupil ellipse for each individual video, it is not compatible with automated large scale HPC or large scale batch processing. In contrast, when deployed on HPC, NeuroPupil achieved further speedups, reducing total processing time by more than an order of magnitude compared to Facemap executed on a local workstation (**Fig. 2H**). This performance advantage enables rapid analysis of large video datasets that would otherwise require prohibitive processing times using conventional pipelines.

Together, these results demonstrate that NeuroPupil simultaneously improves pupil size estimation accuracy and computational efficiency relative to Facemap, enabling scalable, high-throughput pupillometry for large behavioral datasets and translational neuroscience applications.

### NeuroPupil generalizes across heterogeneous human pupillometry datasets while maintaining high precision and scalability

To assess whether the design principles underlying NeuroPupil generalize beyond controlled animal experiments, we extended the framework to large-scale human pupillometry datasets spanning heterogeneous imaging conditions and recording environments (**Fig. 3**). We analyzed two complementary data sources: high-resolution recordings acquired at the Cleveland Clinic Foundation (CCF) and lower-resolution recordings obtained from the publicly available dataset^24^. These datasets differ substantially in optical quality, camera geometry, illumination conditions, and subject behavior, providing a stringent test of robustness and generalizability (**Supp. Fig. 14**).

**Figure 3.**
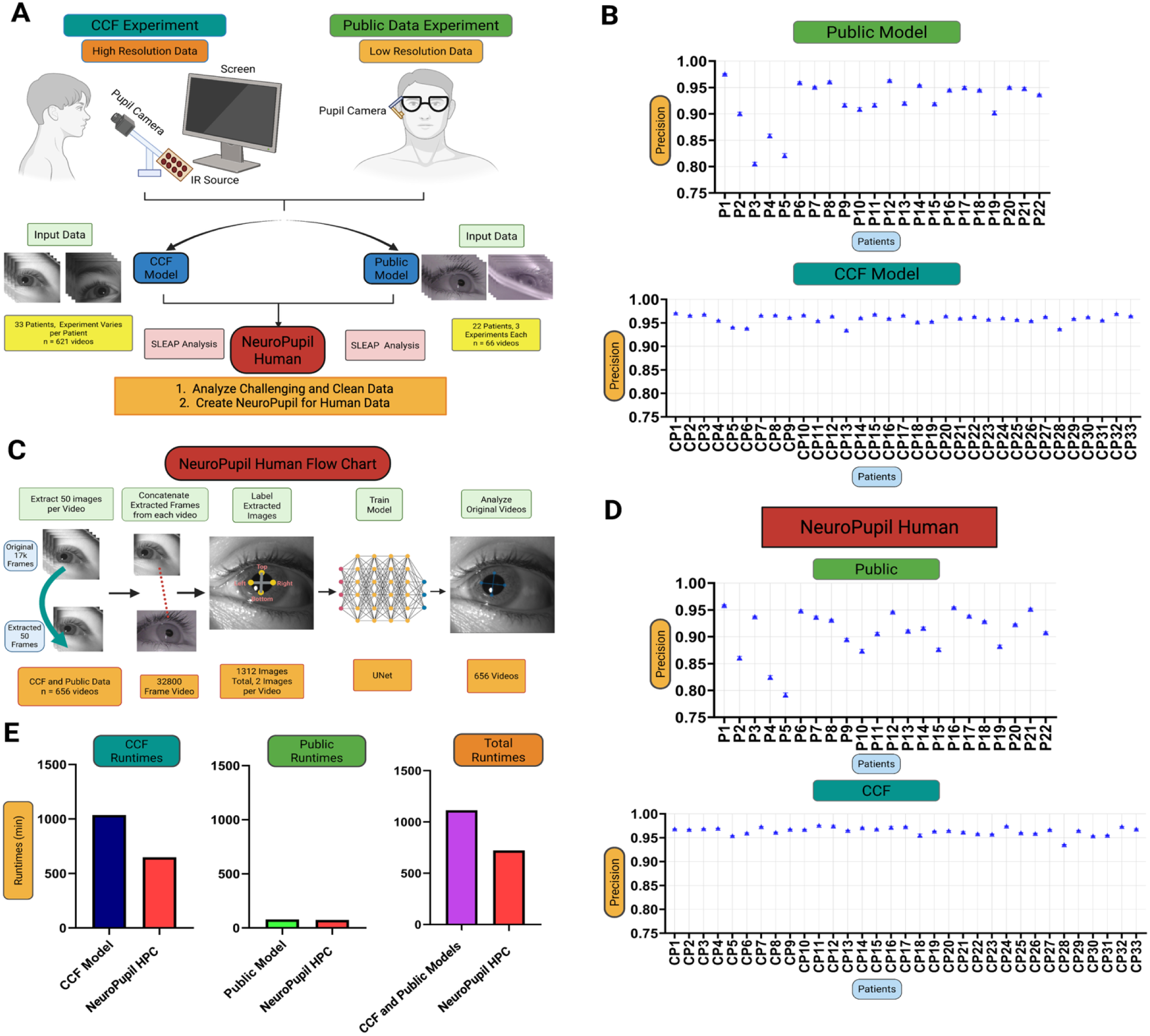
NeuroPupil enables high-precision, scalable pupillometry across heterogeneous human datasets. **(A) Experimental workflow and modeling strategy for human pupillometry across heterogeneous recording conditions**. High-resolution pupil recordings were acquired at the Cleveland Clinic (CCF), while lower-resolution recordings were obtained from a publicly available dataset (citation to be added). Using SLEAP (U-Net backbone) and a pooled multi-subject training strategy analogous to Ten-Mouse Concatenation (TMC), separate models were first trained for the CCF and public datasets. Only two labeled frames per video were required to achieve high precision. Based on this result, a single global model trained on pooled CCF and public data was constructed, termed NeuroPupil Human, to analyze all human pupillometry data. **(B) Precision metrics for dataset-specific models trained on public (low-resolution) and CCF (high-resolution) recordings**. High precision is maintained across subjects despite substantial differences in image quality, resolution, and experimental context. **(C) NeuroPupil Human analysis pipeline**. From each video (n = 656), 50 frames were uniformly sampled and concatenated using a custom graphical user interface. A total of 1,312 images were manually annotated with four pupil boundary keypoints (top, bottom, left, right) and used to train a U-Net–based SLEAP model. The trained NeuroPupil Human model was subsequently applied to full video datasets for automated, frame-by-frame pupil tracking. **(D) Precision of NeuroPupil Human across individual subjects for both public and CCF datasets**. NeuroPupil maintains consistently high accuracy across participants and recording conditions, demonstrating robust generalization across heterogeneous human pupillometry data. **(E) Runtime benchmarking of human pupillometry pipelines**. Processing times are compared for dataset-specific CCF and public models and for NeuroPupil Human deployed on high-performance computing (HPC) infrastructure. NeuroPupil achieves the fastest runtimes, enabling efficient analysis of large-scale human pupillometry datasets.

We first evaluated whether the pooled multi-subject training strategy identified in mice remains optimal for human data. Using SLEAP with a U-Net backbone, we trained multi-subject concatenation models separately for the CCF and public datasets (**Supp. Figs. 15–16**), labeling only two frames per video. Despite this minimal annotation density, both models achieved high precision across subjects within their respective datasets (**Fig. 3B**), demonstrating that sparse labeling combined with pooled training is sufficient for accurate human pupillometry.

Motivated by these results, we trained a single global human model, **NeuroPupil Human**, by pooling data across both CCF and the public dataset (**Fig. 3A**). This model was trained on 1,312 manually annotated images drawn from 656 videos, each labeled with four pupil boundary keypoints (top, bottom, left, right), using a standardized and scalable annotation workflow (**Fig. 3C).** The majority of annotated frames (∼90%) originated from the CCF dataset due to its substantially larger number of videos (n = 590) compared to the public dataset (n = 66), even though the training dataset incorporated using both datasets. Notably, this annotation strategy mirrors that used for mouse data, indicating that NeuroPupil supports a unified, species-agnostic approach to pupillometry.

NeuroPupil Human maintained high precision across individual subjects in both public (**Supp. Fig. 17**) and CCF (**Supp. Fig. 18**) datasets (**Fig. 3D**), despite substantial variability in image resolution, eyelid occlusion, pupil contrast, and experimental context. Importantly, the increased diversity of training data in NeuroPupil Human led to higher precision on the challenging public dataset relative to the public-only model, demonstrating that heterogeneous training enhances generalization while maintaining performance comparable to, or exceeding, dataset-specific models.

In addition to improved accuracy, NeuroPupil Human demonstrated substantial computational advantages. Runtime benchmarking revealed that NeuroPupil Human, when deployed on high-performance computing infrastructure, processed human pupillometry data significantly faster than dataset-specific CCF or public models executed independently (**Fig. 3E; Supp. Fig. 19**). This acceleration enables rapid analysis of hundreds of videos and millions of frames, transforming human pupillometry into a practical, high-throughput measurement suitable for large clinical and translational studies.

Together, these results demonstrate that NeuroPupil extends seamlessly from animal to human pupillometry, maintaining high precision with minimal manual annotation while scaling efficiently to heterogeneous, real-world datasets.

### NeuroPupil outperforms Facemap in accuracy and scalability for human pupillometry under real-world conditions

To directly evaluate NeuroPupil against an established human pupillometry baseline, we compared its accuracy and computational performance with Facemap using heterogeneous datasets derived from both public sources (**Supp. Videos 5-6**) and CCF recordings (**Supp. Videos 7-8**) (**Fig. 4**). These datasets encompass substantial variability in imaging resolution, illumination, eye pose, eyelid occlusion, and motion artifacts, providing a stringent test of performance under real-world conditions.

**Figure 4.**
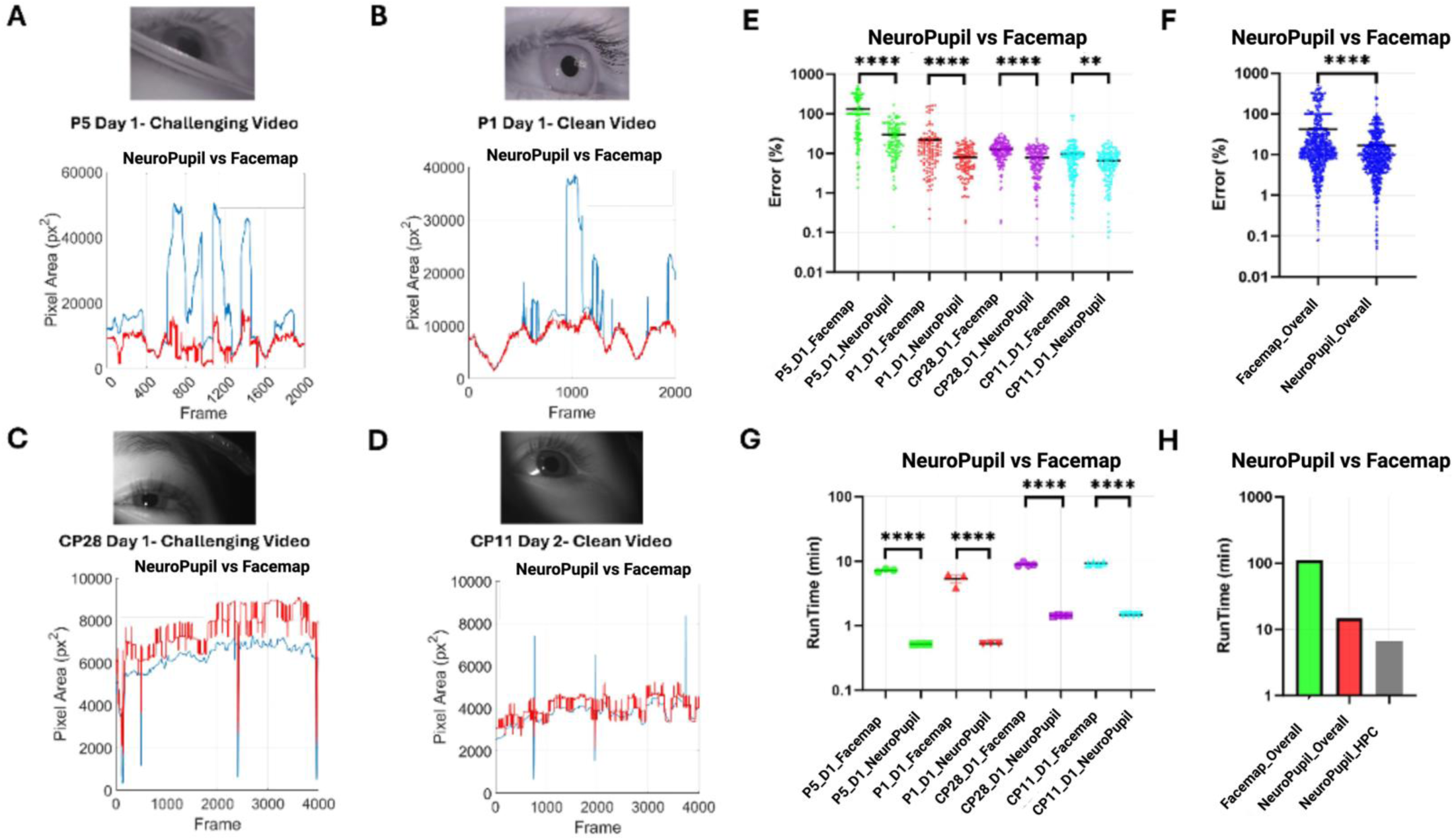
| NeuroPupil outperforms Facemap in accuracy and computational efficiency for human pupillometry. (A–D) Representative pupil area time series comparing Facemap (blue) and NeuroPupil (red) for individual participants across public and Cleveland Clinic (CCF) human datasets: **(A)** P5, day 1; (**B)** P1, day 1; **(C)** CP28, day 1; and **(D)** CP11, day 2. Insets show representative eye images for each recording, highlighting variability in image quality, illumination, and eye pose. **(E) Percent error of pupil area estimates relative to human-annotated ground truth**. Ten frames per video were manually annotated by four independent expert annotators. Error distributions are shown separately for each participant (P5, P1, CP28, CP11), comparing Facemap and NeuroPupil. NeuroPupil exhibits significantly lower error across all participants (two-sided t-test, ****p < 0.0001). **(F) Overall percent error aggregated across all participants and sessions**. NeuroPupil demonstrates significantly improved agreement with human annotations relative to Facemap (two-sided t-test, ****p < 0.0001). **(G) Runtime comparison for individual videos across participants.** NeuroPupil consistently achieves faster processing times than Facemap across all datasets (two-sided t-test, ****p < 0.0001). **(H)** Overall runtime comparison across all videos and participants. Processing times are shown for Facemap executed on a local workstation, NeuroPupil executed on a local workstation, and NeuroPupil deployed on high-performance computing (HPC) infrastructure. NeuroPupil achieves the fastest runtimes, enabling efficient large-scale analysis of human pupillometry data.

Representative pupil area traces illustrate that NeuroPupil closely follows physiologically plausible pupil dynamics while substantially reducing spurious fluctuations and tracking failures observed in Facemap outputs (**Fig. 4A–D**). In particular, Facemap frequently exhibited abrupt excursions and discontinuities in pupil area estimates during partial occlusion, specular reflections, or extreme gaze angles, whereas NeuroPupil maintained stable tracking across frames and sessions.

Quantitative comparison using human-annotated ground truth revealed that NeuroPupil consistently achieved significantly lower error than Facemap across all participants and sessions (**Fig. 4E**). These improvements were observed across both public and CCF datasets, indicating robustness to differences in recording hardware, resolution, and experimental context. When aggregated across participants, NeuroPupil demonstrated a marked reduction in overall error relative to Facemap (**Fig. 4F**; two-sided t-test, ****p < 0.0001).

NeuroPupil also provided substantial gains in computational efficiency. Runtime analyses showed that NeuroPupil processed individual videos significantly faster than Facemap across all participants (**Fig. 4G**). When deployed on high-performance computing infrastructure, NeuroPupil achieved further acceleration, reducing total processing time by more than an order of magnitude relative to Facemap executed on a local workstation (**Fig. 4H**). This performance enables rapid analysis of large-scale human pupillometry datasets that would otherwise be impractical using conventional pipelines.

Together, these results demonstrate that NeuroPupil simultaneously improves agreement with expert human annotations and dramatically accelerates processing for human pupillometry. By combining accuracy, robustness, and scalability, NeuroPupil enables high-throughput analysis of human pupil dynamics in both experimental and clinical research settings.

### NeuroPupil enables functional decoding of brain activity and disease-relevant pupil dynamics

To test whether improvements in tracking precision translate into biologically meaningful signals, we applied NeuroPupil to two downstream analyses: prediction of large-scale cortical activity in behaving mice and classification of disease-associated pupil dynamics in human clinical data (**Fig. 5**). We first examined whether fine-grained pupil dynamics extracted by NeuroPupil encode information about distributed brain activity. GCaMP6s mice underwent six-day experiments combining high-speed pupil videography (40 Hz) with cortex-wide one-photon calcium imaging (20 Hz) during airball locomotion, enabling simultaneous measurement of pupil dynamics and neural activity across ten anatomically defined cortical regions (**Fig. 5A**). Pupil videos were processed using NeuroPupil to extract x/y coordinates of four pupil boundary landmarks (top, bottom, left, right), which were preprocessed and used as inputs to feed-forward neural networks trained to predict regional calcium signals.

**Figure 5.**
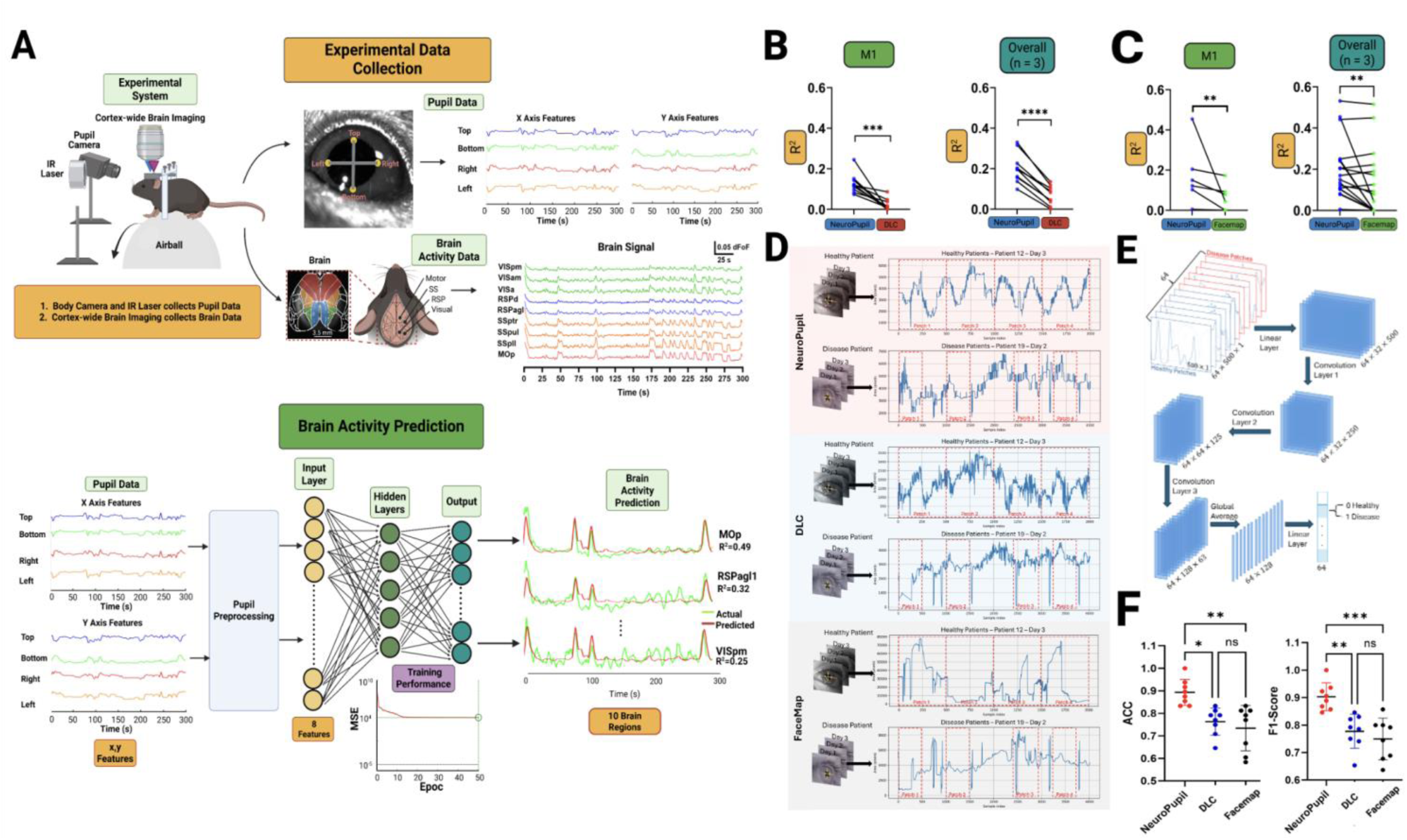
| NeuroPupil enables functional decoding of brain activity and robust classification of disease-related pupil dynamics across species. (A) Experimental setup, data acquisition, and brain activity prediction framework. High-speed pupil videography (40 Hz) was combined with cortex-wide one-photon calcium imaging (20 Hz) in head-fixed GCaMP6s mice running on an airball. Pupil videos were processed using **NeuroPupil** to extract x/y coordinates of four pupil boundary landmarks (top, bottom, left, right). These features were preprocessed and used as inputs to feed-forward neural networks to predict calcium activity across ten anatomically defined cortical regions. Models with varying hidden-layer depths were evaluated. **(B) Comparison of NeuroPupil and DeepLabCut for brain activity prediction using landmark-based features.** Prediction performance was quantified using the coefficient of determination (R²). Top: results from a representative single mouse (M1). Bottom: average performance across all animals (n = 3). Both methods used eight landmark-derived pupil features. NeuroPupil consistently outperforms DeepLabCut at both the single-animal and group levels (***p < 0.001, ****p < 0.0001). **(C) Comparison of NeuroPupil and Facemap for brain activity prediction using pupil area.** Prediction performance (R²) is shown using a single pupil-area feature. Top: representative single mouse (M1). Bottom: average across all animals (n = 3). NeuroPupil achieves significantly higher prediction accuracy than Facemap at both individual and group levels (**p < 0.01). **(D) Human pupillometry examples and patch extraction.** Representative pupil time series from healthy controls and disease patients (strabismus) are shown for three tracking methods (NeuroPupil, DeepLabCut, and Facemap). Red dashed boxes indicate fixed-length temporal patches extracted from continuous recordings for downstream classification. NeuroPupil preserves fine-scale temporal dynamics and signal continuity compared to alternative methods. **(E) 1D convolutional neural network (1D-CNN) architecture for ocular disease classification.** Pupil patches (64 × 500 × 1) from healthy (blue) and disease (red) subjects were used as inputs to a 1D-CNN consisting of stacked convolutional layers, followed by global average pooling and a linear classification layer, yielding a final feature representation of size 1 × 64. **(F) Classification performance across pupil tracking methods.** Left: classification accuracy (ACC). Right: F1-score. NeuroPupil significantly outperforms DeepLabCut and Facemap in distinguishing healthy and disease-associated pupil dynamics (*p < 0.05, **p < 0.01; ns, not significant).

Using an eight-feature landmark-based model, NeuroPupil-derived pupil features enabled significantly higher prediction accuracy than DeepLabCut-derived features, both for a representative single animal (Mouse 1) and when averaged across animals (n = 3) (**Fig. 5B**; **Supp. Fig. 20A**; coefficient of determination, R²; ***p < 0.001, ****p < 0.0001). The eight-feature model successfully predicted 10 out of 10 cortical brain regions, the motor (MOp), somatosensory (SSp_ll, SSp_ul, SSp_tr), retrosplenial (RSPagl, RSPd, RSPv) and visual (VISa,VISam, VISpm) regions, with improvements were consistent across brain regions, indicating that NeuroPupil captures behaviorally relevant pupil dynamics that predict distributed cortical activity.

To enable comparison with Facemap, which outputs only pupil area, we trained an additional single-feature prediction model using pupil area alone. Under the constrained setting, the one-feature model predicted only 6 out of the 10 cortical brain regions (MOp, RSPagl, RSPv1, SSp_ll, SSp_ul, SSp_tr, and VISam), with NeuroPupil-derived pupil area yielded significantly higher prediction accuracy than Facemap, both for a single animal and when averaged across animals (**Fig. 5C**; **Supp. Fig. 20B**; **p < 0.01).

We next evaluated the impact of NeuroPupil on human pupillometry by analyzing disease-associated pupil dynamics in clinical recordings. Representative pupil area time series from healthy controls and patients with strabismus highlight qualitative differences across tracking methods (**Fig. 5D**). We extracted fixed-length temporal patches from these recordings and used as inputs to a one-dimensional convolutional neural network (1D-CNN; **Fig. 5E**). We processed pupil patches (64 × 500 × 1) through stacked convolutional layers followed by global average pooling, yielding compact feature representations for binary classification.

Across classification metrics, NeuroPupil significantly outperformed DeepLabCut and Facemap in distinguishing healthy versus disease-associated pupil dynamics (**Fig. 5F**). NeuroPupil achieved higher classification accuracy and F1-scores (*p < 0.05, **p < 0.01, ***p<0.001), whereas alternative methods showed reduced performance and greater variability. These results demonstrate that small errors in pupil boundary estimation can substantially degrade downstream classification performance and that NeuroPupil preserves diagnostically relevant temporal structure in pupil dynamics.

Together, these analyses demonstrate that NeuroPupil not only improves pupil tracking accuracy but also preserves biologically and clinically meaningful information, as models trained on multiple pupil features capture richer and complimentary dimensions of pupil dynamics than a single-parameter approaches, enabling more accurate and spatially comprehensive prediction of brain-wide neural activity in mice and robust classification of disease-associated pupil dynamics in humans.

## DISCUSSION

Pupillary dynamics are increasingly recognized as a sensitive and quantitative readout of internal brain state, reflecting distributed neural activity, arousal fluctuations, and neuromodulatory tone across species^1–3^. Despite this growing recognition, computational pupillometry has remained methodologically fragmented, with performance often dependent on per-session optimization, extensive manual annotation, and limited scalability. In this study, we introduce NeuroPupil and show that principled, generalization-first design substantially improves the reliability, interpretability, and translational relevance of pupillometry across experimental paradigms, disease models, and species.

A central contribution of NeuroPupil is demonstrating that pooled multi-subject training, when combined with an architecture optimized for spatial continuity (U-Net), yields robust and generalizable pupil tracking across diverse experimental conditions. Through systematic comparative analysis across behavioral contexts, imaging resolutions, and model configurations, we show that data aggregation plays a critical role in performance gains, with substantially greater impact than increased annotation density alone. These findings challenge prevailing pupillometry workflows that rely on per-session or per-animal training^8,15^ and instead motivate generalization as an explicit design objective. By leveraging sparse but diverse training data, NeuroPupil learns representations of pupil geometry that remain stable across motion regimes, optical conditions, and biological variability.

NeuroPupil also addresses key practical limitations of existing pipelines, including sensitivity to video quality, failure modes under extreme dilation or occlusion, and prohibitive computational cost at scale. Although tools such as DeepLabCut^20^ and Facemap^7^ have enabled important advances, our results reveal systematic trade-offs between accuracy, robustness, and throughput when these methods are applied to large, heterogeneous datasets. By integrating optimized training strategies, automated workflows, and compatibility with high-performance computing environments, NeuroPupil achieves lower estimation error and orders-of-magnitude faster processing, enabling pupillometry to scale to millions of frames without manual intervention.

Importantly, improvements in tracking fidelity had direct consequences for downstream biological inference. Using NeuroPupil-derived pupil features, we achieved significantly higher accuracy in predicting distributed cortical activity in behaving mice compared to alternative approaches (**Fig. 5A–C**). These results demonstrate that fine-grained pupil dynamics contain information predictive of brain-wide neural activity and that even small errors in pupil boundary localization can substantially degrade neural decoding performance. Behavioral measurement precision thus emerges as a limiting factor for systems-level neuroscience, rather than a secondary technical concern^15,16^.

Crucially, this biological relevance extends to proof-of-principle disease classification in human data. NeuroPupil preserved fine-scale temporal dynamics in human pupil recordings that enabled modest but significant classification of strabismus compared to DeepLabCut and FaceMap (**Fig. 5D–F**). These results demonstrate that pupil dynamics contain disease-relevant information and that subtle tracking biases can propagate into substantial errors in downstream classification when pupil features are used for analysis. Importantly, these findings should be interpreted as a proof of concept rather than a disease-specific diagnostic claim; larger, more diverse cohorts will be required to evaluate potential clinical utility and generalizability.

A key strength of NeuroPupil is its robustness across neurological disease models and behavioral contexts that introduce substantial variability in pupil dynamics. Stable performance was observed across preclinical models of glioblastoma, PTEN-associated autism spectrum disorder, and cuprizone-induced demyelination^13,14,25^, as well as in human clinical recordings. This robustness supports the suitability of NeuroPupil for longitudinal and disease-focused studies, where behavioral variability is intrinsic rather than confounding.

NeuroPupil was designed with accessibility and reproducibility as core principles. We provide two complementary GUIs that operate sequentially within a unified workflow to replicate High Performance Computing-style analysis workflow, enabling end to end custom NeuroPupil model creation, large-scale inference, and data export for laboratories without access to expensive HPC Infrastructure, while standardized annotation strategies, automated data aggregation, and compatibility with both local and high-performance computing environments lower the barrier to large-scale pupillometry analysis for users with diverse computational expertise. In this framework, increasing dataset size enhances statistical power rather than analytic burden, facilitating reproducible comparisons across laboratories and studies^22,23^.

In summary, NeuroPupil provides a unified and scalable solution for high-precision pupillometry in mice and humans. By demonstrating that benchmarking-driven, generalization-first design improves neural decoding, disease-related classification, and computational efficiency, this work establishes pupillometry as a biologically informative measurement suitable for integrative neuroscience and translational applications.

## RESOURCE AVAILABILITY

### Lead contact

Further information and requests for resources and data should be directed to and will be fulfilled by the lead contact, Murat Yildirim.

### Materials availability

This study did not generate new unique reagents.

### Data and code availability

- The calcium imaging and orofacial datasets presented in this paper are available in the Dryad Repository
- This paper reports original codes and graphical user interfaces that we provided on our github page: (https://github.com/KemalOz21/NeuroPupil)
- Any additional information required to re-analyze the data reported in this paper is available from the lead contact upon request.

## ACKNOWLEDGEMENTS

This work was supported by US National Institute of Health (NIH) grants #R00EB027706 (MY), Case Comprehensive JumpStart Grant #3209 (MY), American Brain Tumor Association Discovery Grant #DG2500080 in memory of Kaitlyn Berg (MY), Cleveland Clinic Research Startup (MY), Case Western University SOURCE Fellowship (KO, FB), National Institutes of Health (NIH) grants (R01 1R01NS112499-01A1 to T.P.L and I.F.M.), Veterans Affair Puget Sound Healthcare System (5I01ABX005978-2 to T.P.L and I.F.M.), ASAP-GP2 (T.P.L and I.F.M.), Michael J. Fox Foundation (MJFF-026283 I.F.M.), American Parkinson Disease Association (APDA) (I.F.M), Parkinson’s Foundation (PDGENE-1333334 I.F.M). Figures 1A, 2A–B, 3A, 4, 5 and Supplementary Figure 1, were created using BioRender.com.

## AUTHOR CONTRIBUTIONS

Conceptualization, K.O. and M.Y.; methodology, K.O., T.C., M.E.L, K.K., B.D., T.P.L., and M.Y.; software, K.O., C.O., T.C., K.K., B.D., T.P.L., and F.B.; validation, K.O., T.C., F.B., and K.K.; formal analysis, K.O., T.C., S.G., K.H., F.B, Z.Z., ; investigation, T.C., M.E.L., M.M., K.O., K.K., F.B., M.Y. ; resources, C.E., N.S., J.D.L., A.S., and M.Y.; data curation, K.O, T.C., F.B., M.M., and M.E.L.; writing – original draft, K.O. and M.Y.; writing – review & editing, K.O., T.C., K.K.,, M.E.L., M.M., F.B., T.P.L., C.O., A.S., N.S., A.C., B.T., C.E., F.G., J.D.L., I.M., and M.Y.; visualization, K.O., T.C., B.D., and M.Y. ; supervision, M.Y.; project administration, M.Y.; funding acquisition, M.Y.

## METHODS

### Animals

All experiments were conducted in accordance with the guidelines of the Cleveland Clinic Institutional Animal Care and Use Committee (IACUC). Mouse genotyping was performed on ear punches by TransnetYX (Cordova, TN) using their automated real-time PCR platform. Mice were maintained on a 14:10 light: dark cycle with ad libitum access to food and water. The room temperature was regulated between 18 °C and 26 °C. Male and female littermates were separated by sex after weaning and housed with same-sex littermates. Reporter (transgenic) mice were generated by crossing *Ai162 (TIT2L-GC6s-ICL-tTA2)-D* (stock no. 031562, Jackson Laboratory) with *B6.Cg-Tg(Camk2a-cre)T29-1Stl/J mice* (CaMKIIα-Cre, stock no. 005359, Jackson Laboratory).

Autism spectrum disorder (ASD)–related phenotypes were studied using the cytoplasmic-predominant *Pten^Y68H/+^* mouse model, originally described previously^26^. This line was backcrossed for >10 generations onto a C57BL/6J background (The Jackson Laboratory, Bar Harbor, ME). For experimental cohorts, male Pten^Y68H/+^ mice on the C57BL/6J background were bred with wild-type female mice (C57BL/6J; The Jackson Laboratory) to generate the mice used in this study.

Multiple sclerosis (MS)–like demyelination was induced by administering cuprizone at a concentration of 0.3% (w/w) to wildtype mice in their diet. Cuprizone-containing food pellets were obtained from Envigo and provided ad libitum for 8 weeks to induce demyelination. Food pellets were replenished by the experimenter three times per week throughout the treatment period. Following the demyelination phase, mice were returned to a standard chow diet for a 6-week remyelination period.

Glioblastoma (GBM) mice were generated by stereotaxic implantation of the SB28 tumor cell line (NRasV12/Pt2/Shp53/mPDGF) acquired from Dr. Hideho Okada at UCSF to wildtype mice. A volume of 3–5 µL of tumor cell suspension was pressure-injected unilaterally into the left dorsolateral striatum (A.P. +1.2 mm, M.L. −1.6 mm, D.V. −3.8 mm relative to bregma) using a 31-gauge needle mounted on a stereotaxic frame (RWD, #68535). Following injection, the craniotomy was sealed with bone wax, and animals were allowed to recover under standard postoperative care.

### Surgical Procedures

Surgical procedures for GCaMP6s, ASD and MS mice closely followed the procedure previously used for whole-cortex widefield imaging^27^. Mice were anesthetized with isoflurane (induction 2.5%; maintenance 1-1.5%). Buprenorphine (3.25 mg/kg), Meloxicam (5 mg/kg), and sterile saline (0.05 mL/g) were administered at the start of surgery. Anesthesia depth was confirmed by toe pinch. Hair was removed from the dorsal scalp (Nair, Vetiva Mini Hair Trimmer) and the area was disinfected with 3 alternating applications of betadine and 70% isopropanol. Bupivacaine (5 mg/kg) was then injected under the skin for local anesthetic before the scalp was removed. The skull was then exposed, cleaned, and dried. The remaining outer skin was affixed in position with tissue adhesive (Vetbond, 3M) for clean surgical margins. We created an outer wall using dental cement (C&B Metabond, Parkell) while leaving as much of the skull exposed as possible. A custom circular headbar (eMachineShop) was secured in place using dental cement (C&B Metabond, Parkell). A layer of optical glue (Norland Optical Adhesive NOA 81, Norland Products) was then applied to the exposed skull and cured with a UV flashlight (LIGHTFE, UV301Plus-365nm). The mice were allowed to fully recover in a warmed chamber and then returned to their home cages. Post-operational care consisted of three daily injections of Meloxicam (5 mg/kg) following the surgery.

In addition to the surgical procedures described above, glioblastoma (GBM) mice underwent stereotaxic implantation of the SB28 tumor cell line during the same surgical session. Following skull exposure, a small craniotomy was made above the left dorsolateral striatum, and 3–5 µL of SB28 tumor cell suspension was pressure-injected unilaterally at A.P. +1.2 mm, M.L. −1.6 mm, D.V. −3.8 mm relative to bregma using a 31-gauge needle mounted on a stereotaxic frame (RWD, #68535). The craniotomy was sealed with bone wax, after which the procedures for skull preparation, headbar fixation, optical clearing, and wound closure were completed as described above. Postoperative recovery and analgesic care were identical to those described for other experimental groups.

### Data Acquisition, Triggering, and Synchronization

Animal locomotion was measured using a custom air-ball system that allowed free planar movement. The airball system was composed of a 20-cm polystyrene foam ball and a hemispherical 3D printed PLA bowl with an internal diameter that fitted with the ball modified from previous studies^28,29^. The bowl had eight holes for pressured air at the bottom. The head of the mouse was fixed to a rigid head mount bar and posts via the head plate and positioned ∼1 cm above the top of the ball. Ball motion was tracked using two orthogonally mounted USB optical mice (Gaming Mouse G302, Logitech), capturing forward–backward, lateral, and rotational components of movement. Raw motion signals were read by a Raspberry Pi, converted into three analog voltages via a digital-to-analog converter, and streamed to a data acquisition (DAQ) system (National Instruments PCI-6321 and BNC-2110).

Widefield brain imaging and orofacial video acquisition were performed using cameras operating in hardware-triggered mode. Each camera (Thorlabs, CS505MU1, Kiralux) was controlled by an Arduino microcontroller that initiated frame acquisition only when receiving a synchronization signal from the DAQ system. This ensured precise temporal alignment across imaging modalities. Brain imaging used alternating blue and violet illumination, implemented by the Arduino at 40 Hz, with successive frames captured under alternating wavelengths to enable spectrally distinct measurements.

### Widefield Imaging of Brain Activity

For wide-field imaging, cortical excitation was performed sequentially using blue (M470L5, Thorlabs) and violet (M405LP1, Thorlabs) light emitting diodes (LEDs). Blue illumination (470 nm) targeted genetically-encoded calcium indicators (GECI) (GCaMP6s), while violet (405 nm) illumination immediately followed to provide a reference measurement used to correct hemodynamic artifacts. The two excitation wavelengths were combined through a dichroic beamsplitter (87-063, Edmund Optics) and sent onto the cortical surface via an objective lens (MVL50M23, Navitar). Emitted light was collected through an emission dichroic beamsplitter (T495LPXR, Chroma), and filtered using a band-pass filter (86-963, Edmund Optics). The filtered fluorescence signal was then focused on a CMOS camera (CS505MU1, Kiralux, Thorlabs) through a second objective lens (MVL16M23, Navitar). Data was acquired at a resolution of 1200 × 1200 pixels with 4×4 pixels binning, resulting in an effective 300×300 pixels covering a cortical area of 9.85 × 9.85 mm^2^. This configuration enabled an acquisition rate of 40 frames per second (fps), sufficient for resolving the kinetics of the GCaMP6s calcium indicator. Experiments were carried out at 40 Hz frame rate for blue and violet light combined (effective frame rate of 20 Hz). Average laser powers were maintained below 10mW for the blue LED and below 50 mW for violet LED. Beam diameters were measured by the knife-edge test method, (1/e^2^ criterion) were 9.2 for blue LED and 11.7mm for violet LED.

### Allen Brain Atlas Registration

We used the alignment script provided by the locaNMF toolbox^30^ to rigidly align the skull surface’s anatomical landmarks to a standard template atlas. For each animal, we selected seven key points: the left, center, and right intersections between the anterior cortex and olfactory bulb, the location of the bregma, and the base of the retrosplenial cortex on the midline skull suture.

### Hemodynamic Subtraction

For hemodynamic correction, we first computed the blue and violet ΔF/F for each brain region by subtracting and dividing by the median average brain activity during the whole session (5 min). We used linear regression to regress the violet time series on the blue time series, i.e. find coefficients m and b of the regression b = mv + b where b is the blue and v is the violet ΔF/F signal. The subtracted signal was given by b – (mb + b). This gives a single time series per region, which was used for subsequent analyses.

### Imaging of Pupil Features

We recorded the pupil features of the animal in all mouse lines. To illuminate the left side of the face, we used an infrared light source (LIU780A, Thorlabs). The pupil images were collected with a focusing telecentric lens (58-430, Edmund Optics) and a monochromatic camera (1500M-GE-TE, Thorlabs) for the wheel experiments, and another camera (CS505MU1, Thorlabs) with the same telecentric lens for the airball experiments. Both cameras acquired the pupil data with the frame rate of 40 Hz. We utilized 1200×1200 pixels with binning 4×4pixels (effectively 300×300 pixels) which cover 4.11 × 4.11 mm^2^ surface area on the mouse eye.

### Wheel Data Acquisition

Animal were trained on a version of dynamic foraging paradigm with an identical procedure as described previously^31,32^. Briefly, the training apparatus and software for the training were adapted from the Rigbox framework for psychophysics experiments in rodents^33^. Mice were head-fixed on the platform (built from Thorlabs hardware parts) and their body placed in a polypropylene tube to limit the amount of movement and increase comfort. Their paws rested on a vertical Lego wheel (radius 31 mm) which was coupled to a rotary encoder (E6B2-CWZ6C, Omron), which provided input to a data acquisition board (BNC-2110, National Instruments). The data acquisition board also provided outputs to a solenoid valve (#003-0137-900, Parker) which controlled the water reward delivery.

Widefield brain imaging and pupil video acquisition were performed using cameras operating in hardware-triggered mode. Each camera (brain imaging-Thorlabs, CS505MU1, Kiralux; pupil imaging-1500M-GE-TE, Thorlabs) was controlled by an Arduino microcontroller that initiated frame acquisition only when receiving a synchronization signal from the DAQ system. This ensured precise temporal alignment across imaging modalities. Brain imaging used alternating blue and violet illumination, implemented by the Arduino at 40 Hz, with successive frames captured under alternating wavelengths to enable spectrally distinct measurements.

### Airball Data Acquisition

Animal locomotion was measured using a custom air-ball system that allowed free planar movement. The airball system was composed of a 20-cm polystyrene foam ball and a hemispherical 3D printed PLA bowl with an internal diameter that fitted with the ball modified from previous studies^28,29^. The bowl had eight holes for pressured air at the bottom. The head of the mouse was fixed to a rigid head mount bar and posts via the head plate and positioned ∼1 cm above the top of the ball. Ball motion was tracked using two orthogonally mounted USB optical mice (Gaming Mouse G302, Logitech), capturing forward–backward, lateral, and rotational components of movement. Raw motion signals were read by a Raspberry Pi, converted into three analog voltages via a digital-to-analog converter, and streamed to a data acquisition (DAQ) system (National Instruments PCI-6321 and BNC-2110).

Widefield brain imaging and pupil video acquisition were performed using cameras operating in hardware-triggered mode. Each camera (Thorlabs, CS505MU1, Kiralux) was controlled by an Arduino microcontroller that initiated frame acquisition only when receiving a synchronization signal from the DAQ system. This ensured precise temporal alignment across imaging modalities. Brain imaging used alternating blue and violet illumination, implemented by the Arduino at 40 Hz, with successive frames captured under alternating wavelengths to enable spectrally distinct measurements.

### Public Human Data Acquisition

We evaluated NeuroPupil on a publicly available human pupillometry dataset, Labelled Pupils in the Wild (LPW), which contains eye-region videos recorded under unconstrained, real-world conditions^24^. The dataset includes recordings from 22 participants, each contributing three eye videos, yielding a total of 66 videos. Videos were acquired using a head-mounted dark-pupil eye tracker at approximately 95 frames per second with a spatial resolution of 640 × 480 pixels, resulting in 130,856 frames overall. Recordings span diverse indoor and outdoor environments, illumination conditions, gaze directions, and participant characteristics, including the presence of glasses, contact lenses, and eye makeup. All frames are accompanied by manually verified ground-truth pupil annotations. For our analyses, we used the raw eye videos and associated annotations as provided, without modification, and treated each participant as an independent unit to avoid subject-level information leakage.

### CCF Human Data Acquisition

Human eye-movement and pupillometry data were acquired in the Ocular Motility Laboratory directed by Dr. Fatema Ghasia at the Cole Eye Institute. Experiments were conducted in dedicated luminance-controlled rooms with independently dimmable lighting to ensure stable and reproducible visual conditions. Eye movements and pupil dynamics were recorded using clinically validated infrared video eye-tracking systems, including EyeLink 1000 and EyeLink 1000+ platforms (SR Research), supporting binocular pupil tracking with sampling rates of up to 1000–2000 Hz in both remote and mounted configurations. Additional recordings were obtained using complementary systems for non-invasive eye and head tracking, including EyeLink II, portable head–eye trackers, and wearable eye-tracking platforms, enabling flexibility across patient age groups and experimental paradigms.

Visual stimuli were presented using calibrated displays and immersive systems as appropriate, including high-resolution LCD monitors, tablet-based platforms with integrated head and eye tracking, and virtual reality headsets for naturalistic viewing conditions. All data were collected under institutional review board approval with informed consent, and recordings were obtained from a clinical cohort of 33 patients, each contributing a variable number of pupil videos across sessions. Raw eye videos and associated tracking outputs were processed uniformly using the same preprocessing and analysis pipeline, and subject identity was preserved throughout analysis to enable patient-level evaluation while preventing information leakage across recordings.

### PTEN Mouse Data Acquisition

Heterozygous Pten^Y68H/+^ mice were used to assess pupil dynamics, locomotion, and large-scale brain activity during controlled behavior. Animals underwent a three-day acclimation protocol to head fixation and the experimental environment. On the first day, mice were head-fixed and allowed to move freely on a foam ball with pressurized air turned off. On the second and third days, pressurized air was gradually introduced and increased to approximately one-quarter and one-half of the final operating pressure, respectively, with the final pressure set to 80 psi.

Following acclimation, seven mice were each recorded across six experimental sessions, during which pupil dynamics, and locomotion were acquired simultaneously (**Supp. Fig. 10**).

### MS Mouse Data Acquisition

Prior to cuprizone exposure, mice underwent the same three-day head-fixation acclimation protocol used for Pten^Y68H/+^ experiments. Animals were then recorded longitudinally during a 5-week baseline period, during which pupil dynamics, and locomotion were acquired repeatedly. Cuprizone treatment was initiated after baseline recordings and continued for 8 weeks, with multimodal recordings maintained throughout treatment. In the present study, analyses focus on pupil measurements acquired during the 5-week baseline period and the first 5 weeks of cuprizone treatment for seven mice (**Supp. Fig. 11**).

### GBM Mouse Data Acquisition

A syngeneic glioblastoma (GBM) mouse model was established using intracranial injection of SB28 tumor cells. Mice underwent the same three-day head-fixation acclimation protocol described for Pten^Y68H/+^, and cuprizone experiments prior to data collection. Following acclimation, three baseline recording sessions were acquired for each animal, during which pupil dynamics and locomotion were recorded prior to tumor implantation.

After SB28 injection, animals were recorded longitudinally with the same acquisition paradigm until reaching predefined humane endpoints. In the present study, analyses focus on three pre-injection baseline sessions and three post-injection sessions acquired on days 1, 3, and 5 following tumor implantations for five mice.

### DeepLabCut-Based Pupillometry Analysis for Animal Videos

DeepLabCut (DLC) version 2.3.10 was utilized as the initial software package to evaluate pupillometry data. DLC offers both command-line and graphical user interface (GUI) functionalities, facilitating its accessibility and ease of use. The initial data analysis approach employed was Single Day Analysis (SDA) (**Supp. Fig. 1a**), wherein a separate model was trained and evaluated for each individual video. Given the fixed camera angle during data acquisition and minimal occlusions within the videos, SDA provided a precise method for individual video evaluation.

Model training for SDA was performed using DLC’s Residual Neural Network (ResNet-50) architecture. Raw videos were first imported into DLC, followed by modification of the configuration file to define the project parameters, including the project name and the number of labeled points. Image extraction was then performed, utilizing either of DLC’s two unsupervised extraction methods, K-means clustering or uniform frame sampling, or a supervised manual extraction method.

K-means clustering grouped video frames based on visual appearance and selected 20 representative frames across different clusters for labeling. Alternatively, uniform sampling selected 20 frames randomly distributed across the video’s spatial and temporal dimensions. Following frame extraction, the pupil’s top, bottom, left, and right edges were manually labeled (**Supp. Fig. 1a**), with the label size adjusted from the default value of 12 to 8 to enhance precision.

Training parameters were standardized across models: shuffle was set to 1, network architecture to ResNet-50, and data augmentation to imgaug. Training was performed on CPU, with a maximum of 1000 iterations, snapshots saved every 500 iterations, an iteration display increment of 50, and a total of five snapshots recorded.

Model performance was subsequently evaluated using DLC’s “Plot Predictions” and “Compare Body Parts” functions. To ensure correct label display in the Analysis tab, the GUI was closed and reopened after model training. Prior to analysis, the appropriate raw video was loaded, and options for saving results as CSV, plotting trajectories, and (optionally) filtering predictions were available; however, filtering was not applied in this study.

Following analysis, labeled videos were generated by selecting the corresponding analyzed video, enabling the “Plot All Body Parts” and “Draw Skeleton” options, and exporting the visualization for qualitative assessment. To verify the robustness of the unsupervised extraction approaches, the model yielding the highest precision for each subject in the wheel dataset was selected for downstream analysis.

To assess the effectiveness of unsupervised frame extraction, we trained additional models on datasets manually extracted with 10 to 100 frames (**Supp. Fig. 2**). For each animal, a single video was selected, from which SDA models were created using images ranging from 10 to 100 frames. Model performance did not differ significantly between those trained on manually extracted frames and those generated using DLC’s unsupervised clustering methods (K-means and uniform sampling) (**Supp. Fig. 3**). Given the limited variability in visual content across frames in the SDA approach, extensive frame extraction was deemed unnecessary. Therefore, DLC’s default unsupervised clustering methods were used for subsequent analyses of the wheel and airball datasets.

The second data training approach utilized was One Mouse Concatenation (OMC) (**Supp. Fig 1B**), which involved constructing and analyzing a single model for each individual animal’s pupil dataset. For each animal, 3000 frames were evenly selected from the raw video recordings, and they were concatenated together into a single video file representing the animal’s full dataset using a Python-based GUI called mediaGUI. Model training for OMC followed the same procedures as in Single Day Analysis (SDA), including manual labeling, configuration of label names in the configuration file, selection of an appropriate image extraction method (unsupervised K-means clustering or uniform sampling), and training of the network using DLC’s default parameters (**Fig. 1A-B**).

Following model training, each original video corresponding to the trained animal model was analyzed individually. A total of twenty OMC models were developed to reflect the pupil data for the twenty animals in the study (ten for wheel and ten for airball experiments). The primary difference between OMC and SDA workflows was that, in OMC, the concatenated video was used for training, whereas in SDA, training was performed on individual experimental videos.

During OMC training, it was observed that DLC’s unsupervised image extraction algorithms did not evenly sample frames across the concatenated videos. For instance, in mouse F11 from the wheel dataset, three extracted frames originated from Day 2, while no frames were selected from Day 7. To ensure balanced temporal coverage across the videos, two frames were manually extracted per day, resulting in 20 manually selected images per animal for model training and analysis (**Fig 1A-B**).

The third data training approach was Ten Mouse Concatenation (TMC), in which a single model was trained and evaluated using data combined across all animals within each dataset type (wheel or airball). For TMC training (**Supp. Fig. 1C**), 300 frames were extracted evenly from each animal experimental video, and concatenated together to create a single TMC video that can analyze all the data in either wheel or airball experiment (**Fig. 1B**). Model labeling, network training, and evaluation procedures were identical to those used in the OMC approach. However, unlike OMC, where models were specific to individual animals, the TMC model represented the full set of videos from the specific experiment. The time taken to create the models (SDA, OMC, and TMC) and analyze them were recorded as well (**Fig. 1G**).

The TMC and OMC analysis workflows followed similar procedures, with the key difference being that TMC evaluated all videos collectively, while OMC assessed each animal’s videos individually. To improve model generalization and performance across the combined datasets, an extra training set was created by manually extracting two frames per individual video, resulting in 200 images in total. The focus was placed on the low-resolution wheel experimental data because the TMC models performed poorly in this context (**Fig. 1B**, **top**), and it was hypothesized that labeling more images might improve precision. In addition to the 200-image training set, multiple TMC models were generated with 100, 400, 800, and 1,600 labeled images, which were then applied to both a particularly challenging wheel experiment dataset (W9) and clean videos (W3) (**Supp. Fig. 7**).

### SLEAP-Based Pupillometry Analysis for Animal Videos

SLEAP version 1.4.1 was utilized as the second software package for training and evaluating pupil video data. Like DeepLabCut (DLC), SLEAP offers both command-line functionality and an integrated graphical user interface (GUI). However, due to the greater number of adjustable parameters, SLEAP’s GUI was found to be significantly more complex to operate compared to DLC. To determine the optimal settings for training and analysis, several SLEAP parameters were systematically tested.

An initial evaluation was performed using a single mouse concatenated video for subject F11 from the wheel dataset (**Supp. Fig. 3**). This video was selected because it included frames from Day 10, which exhibited the lowest tracking precision and the greatest number of occlusions. The goal was to determine the optimal training conditions for challenging video data.

Three different K-means clustering configurations were evaluated, corresponding to 1, 5, and 10 clusters. To maintain consistency with DLC’s unsupervised extraction methodology, the total number of training frames was standardized at 20 images across all clustering conditions. For instance, when using 5 clusters, 4 frames were randomly selected from each cluster, totaling 20 images. In addition, two training frame size settings (32 and 64) were tested to evaluate the effect of input resolution on model performance.

All models were trained using the U-Net neural network architecture. Six combinations of K-means clustering number and frame size parameters were systematically assessed. Analysis was performed using SLEAP’s default post-processing settings: Simple matching combined with Intersection Over Union (IOU) and Hungarian algorithm assignment.

The evaluation revealed that training with 20 images extracted from a single cluster and using a training frame size of 64 yielded the highest precision (**Supp. Fig. 3A**). Based on these findings, subsequent model training procedures employed even frame selection from the concatenated videos, rather than a clustered selection, to optimize model performance.

After determining the optimal training parameters, the same video used for parameter optimization (subject F11, Day 10) was utilized to evaluate tracking strategies (**Supp. Fig. 3B**). Two primary analysis modes were tested in the SLEAP code package: Simple tracking and Optical Flow–based tracking. For each mode, six built-in association strategies were evaluated, consisting of all combinations of three instance-matching metrics, Intersection Over Union (IOU), Centroid Matching, and instance matching, paired with two frame-to-frame assignment algorithms, Greedy and Hungarian, resulting in a total of twelve inference configurations. IOU, commonly used in object detection, measures the overlap between predicted and ground-truth shapes. Centroid matching computes the Euclidean distance between the centroids of predicted and actual labels, while instance matching calculates normalized distances between sets of labels across frames. For assignment, the Greedy method selects the nearest prediction on a frame-by-frame basis, whereas the Hungarian algorithm optimizes global matching efficiency across frames. For each combination of tracking method, instance matching strategy, and assignment algorithm, precision was recorded, resulting in a total of 12 systematically tested parameter combinations (**Supp. Fig. 3B**). The Flow tracking, Centroid matching, and Greedy assignment combination yielded the highest tracking precision.

Following optimization of the training and analysis parameters, these settings were applied to perform Single Day Analysis (SDA), One Mouse Concatenation (OMC), and Ten Mouse Concatenation (TMC) workflows in SLEAP (**Fig. 1A-B, and Supp. Fig. 4,5, and 6**). Each methodology for both the wheel and airball datasets utilizes the optimal parameters derived for model labeling, training, and analysis.

To perform Single Day Analysis (SDA) in SLEAP (**Supp. Fig. 1A**), each raw video was first imported into the SLEAP GUI as a grayscale file. The desired nodes (label points) and their relationships were defined, specifically linking Top with Bottom and Right with Left. Frame selection for labeling was performed using the sample+stride method to ensure that frames were evenly sampled across the duration of each video.

Nodes were manually placed on the designated anatomical landmarks, and the model was trained and evaluated using the optimized parameters identified previously. Prior to model training, the Max Stride and Filters parameters were both set to 64 to match the model’s frame size of 64 pixels. Before analyzing each video and generating labeled visualization videos, the correct node indices corresponding to the labeled points (0, 1, 2, and 3 for four labeled nodes) were specified. Additionally, the optimized Flow tracking, Centroid matching, and Greedy assignment parameters were inserted to ensure consistency during analysis.

For video generation, the frame rate was set to match that of the raw video: 35 frames per second (fps) for the wheel dataset and 40 fps for the airball dataset. After analysis, the CSV output file for each video was saved to enable downstream extraction of each frame’s tracking precision and the x, y coordinates of each labeled node.

For SDA and OMC workflows (**Supp. Fig. 1A-B**), 20 frames were evenly selected and labeled for model training, consistent with the frame count used in DLC’s corresponding methodologies (**Fig. 1A-B**, **Supp. Fig. 4-6**). For TMC, 200 frames were extracted evenly across all videos, matching the number of frames used in DLC’s TMC manual extraction process (**Supp. Fig. 1C**) (**Fig. 1B and Supp. Fig 6**). Each trained model was subsequently analyzed on the respective animal datasets following the same workflow previously established for DLC. The same method used in DLC to evaluate whether precision increases with the number of labeled images was also applied in SLEAP. In addition to the 200-image training set (two images per video), multiple TMC models were generated with 100, 400, 800, and 1,600 labeled images, and applied to a challenging experiment dataset (W9) and clean videos (W3) in the wheel dataset (**Supp. Fig. 7**) because the overall data was low resolution in comparison to the airball data. This analysis showed that precision gains plateaued after two images per video (200 images), with only minimal improvement observed beyond this point. The time taken to create the models (SDA, OMC, and TMC) and analyze them were recorded as well (**Fig. 1G**).

### NeuroPupil Animal Analysis Method

For the Animal NeuroPupil analysis, a global SLEAP model was developed to analyze all animal dataset videos across different experimental conditions (**Fig. 1C and Supp. Fig. 1D**). To construct this model, 150 images were extracted from each video (n = 200) from the low-resolution wheel (n = 100) and high-resolution airball (n = 100) experiments respectively using the **mediaGUI** (**Supp. Fig. 13 and Supp. Fig. 8**).

From this combined dataset, two frames per video were extracted and labeled, resulting in a total of 400 labeled frames across 200 videos. The global model was then trained and analyzed using the same optimized parameters previously established for other SLEAP models, ensuring methodological consistency across experiments.

In addition to analyzing the wheel and airball datasets used during model training, the global model was also applied to independent wheel and airball datasets that were not included in the training set (**Fig. 1D-E and Supp. 9**), allowing assessment of the model’s generalization capabilities. Furthermore, the model was extended to analyze pupil tracking in PTEN (**Fig 1F and Supp. Fig. 10**), Cuprizone (**Fig 1F and Supp. Fig. 11**), and Glioblastoma **(Fig 1F and Supp. Fig. 12**) animal datasets, demonstrating its applicability across multiple experimental paradigms.

Runtimes for the Animal NeuroPupil model were measured on both the local computer (in parallel with the SLEAP TMC analysis) and the HPC system (**Fig. 1G**). Specifically, the time required to create, label, train, and analyze the model for the wheel and airball datasets was recorded (**Fig. 1G**). Following the analysis, two custom-built GUIs were developed to ensure reproducibility for users wishing to create their own NeuroPupil models. The first, **mediaGUI**, facilitates image extraction for model training, while the second, **sleapGUI Pupil**, enables video analysis within the SLEAP 1.4.1 environment. The sleapGUI Pupil allows users to efficiently analyze pupillary data, export results as CSV files, and generate labeled videos using the same parameters applied in our pupil data analyses (**Supp. Fig. 13A**). For users with more technical expertise and want to utilize HPC, the complete workflow for creating NeuroPupil models and analyzing video data on the HPC system is provided (**Supp. Fig. 13B**).

### NeuroPupil Pipeline Architecture and Availability

To facilitate reproducible and accessible analysis of pupillary data, we developed the NeuroPupil pipeline as a modular, open-source framework composed of two complementary graphical user interface (GUI) tools: mediaGUI for video preprocessing and sleapGUI for large-scale model inference and analysis. These tools are designed to be used sequentially within a unified workflow.

To simplify adoption of the pipeline, we created a central GitHub repository, NeuroPupil (https://github.com/KemalOz21/NeuroPupil), which serves as the main entry point for users and provides direct links to the mediaGUI and sleapGUI repositories for installation. This repository includes clear installation instructions for both tools, along with tutorial videos demonstrating how to build and deploy custom NeuroPupil models tailored to different experimental paradigms. The tutorials present a step-by-step “recipe” guiding users from raw video collection, through preprocessing and model training, to efficient large-scale inference and downstream analysis. This structure enables new users to adopt the pipeline with minimal technical expertise while allowing advanced users to adapt the workflow to their own datasets.

The two GitHub repositories correspond to distinct components of the NeuroPupil pipeline that address different computational bottlenecks. mediaGUI standardizes and stabilizes training data preparation, while sleapGUI enables efficient batch analysis and model deployment at scale. Together, they form an integrated pipeline for NeuroPupil -based pupillary tracking. If users encounter issues, unexpected behavior, or have suggestions for improvement, they are encouraged to report them through the issue tracker on the NeuroPupil GitHub page. Reported concerns will be reviewed, investigated, and addressed to ensure continued usability and long-term maintainability of the pipeline.

### mediaGUI Methodology

mediaGUI was developed to efficiently preprocess pupillary video data by extracting and concatenating frames across multiple experimental sessions (**Supp. Fig. 13 and Supp. Fig19**). Users upload raw videos, specify the number of frames to be evenly extracted from each video, and generate a single concatenated output video for model training. Through systematic testing, we found that training stability is optimal when concatenated videos contain ≤30,000 frames at 400 × 400 pixel resolution. At frame counts ≥50,000, model training becomes unstable and memory requirements exceed what is feasible on standard desktop or laptop computers. By standardizing preprocessing, mediaGUI ensures consistent and stable inputs for models trained in SLEAP and can also be used with related frameworks such as DeepLabCut. The tool supports output in both MP4 and AVI formats and minimizes manual effort through a streamlined, minimalistic interface.

mediaGUI is implemented in Python and uses PyQt6 (v6.6.0) for GUI development. Video processing operations are powered by OpenCV-Python (v4.8.1) and NumPy (v1.26.0). Where available, the CUDA Toolkit is used to offload frame reading and resizing to the GPU, significantly accelerating preprocessing. On Windows systems, the openh264 library is integrated to improve MP4 rendering speed relative to AVI output. The software is cross-platform and has been tested on Windows 10/11, macOS 14/15, and Ubuntu 22.04. The software is open-source and available under the MIT License in the main NeuroPupil Github Page. Installation is recommended via a standalone executable but is also supported through pip, with detailed usage instructions provided in the repository.

### sleapGUI Pupil GUI

sleapGUI was developed within the SLEAP environment (**Supp. Fig. 13A and Supp. Fig. 19A**) to automate NeuroPupil model deployment and overcome limitations of the default SLEAP graphical user interface. In the default SLEAP GUI, users are required to manually specify analysis parameters for each individual video, a process that must be repeated for every analysis run and become increasingly time-consuming when processing large datasets. sleapGUI automates this workflow, enabling batch analysis of multiple videos with a single user action. Optimized analysis parameters for large-scale mouse pupillary tracking are embedded directly within the sleapGUI backend commands. Specifically, inference is performed using a predefined combination of Flow, Centroid, and Greedy tracking methods, which we empirically determined to be optimal for NeuroPupil-based pupillary analysis. Because these parameters are hard-coded into the analysis backend, users do not need to manually select or tune inference settings, eliminating repetitive parameter entry and reducing the risk of inconsistent analysis configurations across experiments.

Two sleapGUI variants were developed to support different experimental paradigms. The sleapGUI Pupil is designed for single-animal pupil tracking using the 4-keypoint model and was used to analyze all experimental data in this study except for the social interaction experiments. The sleapGUI variants are designed to mimic HPC batch workflows on local machines, allowing multiple videos to be analyzed sequentially without user supervision. Output files include .slp tracking files, .csv data tables, and labeled visualization videos, which are immediately suitable for downstream analyses such as brain activity prediction.

The applications are implemented in Python and leverage QtPy (v2.4.2), which is pre-installed within the SLEAP environment, for GUI development. Core functionality relies on NeuroPupil command-line interfaces, with the GUI acting as an abstraction layer. Threading is used to maintain interface responsiveness during computationally intensive tasks, and platform-specific optimizations ensure reliable execution on Windows, macOS, and Linux systems. sleapGUI is compatible with SLEAP versions up to v1.4.1. The sleapGUI requires an NVIDIA GPU to analyze the videos in a fast manner. The main NeuroPupil GitHub page provides open-source access to sleapGUI along with detailed installation instructions and usage documentation, including resources to download and install SLEAP v1.4.1, enabling users to run the custom sleapGUI within a supported SLEAP environment.

### SDA, OMC, TMC, and NeuroPupil Mouse Processing Times Methodology

Following data evaluation using DLC and SLEAP, we sought to compare the processing times associated for each model method (SDA, OMC, TMC, and NeuroPupil) with using the Graphical User Interface (GUI) for each software package. In DLC, execution times were compared between the CPU and GPU configurations (**Fig. 1G**). In SLEAP, execution times were compared between the GUI using only the GPU, as SLEAP requires GPU acceleration for model training and analysis. All experiments were conducted on a PC equipped with a 12th Generation Intel(R) Core(TM) i7-12700K processor, 32 GB of RAM, and an NVIDIA GeForce RTX 3080 GPU. For DLC GUI execution in the SDA, OMC, and TMC models, the DLC environment was first activated through the Anaconda Prompt to open the DLC GUI, and the commands were obtained from DLC’s official GitHub documentation.

The SLEAP environment was similarly activated via Anaconda Prompt to activate its GUI and run the SDA, OMC and TMC environments. The commands to open the SLEAP default GUI were obtained from DLC’s official GitHub documentation. NeuroPupil was run inside the SLEAP module on the High-Performance Computer (HPC), using command lines adapted from the SLEAP GitHub page. The commands to analyze the video data were first compiled in a Notepad file and then copied into a batch SLURM script for execution. The challenges associated with manually managing SLEAP command-line workflows, combined with the complexity of scaling analyses for HPC environments, inspired the development of sleapGUI Pupil. This tool was designed to automate video analysis workflows within the SLEAP environment, enabling more efficient and accessible processing for users without requiring extensive command-line expertise.

### High Performance Computing Analysis

HPC was utilized for Neuropupil mouse analysis to process all videos in a high-throughput manner. This was achieved by running a single SLURM job, which leveraged parallel computing to accelerate inference speed (**Supp. Fig. 13B and Supp. Fig. 19B**). A batch SLURM script was created and submitted to the HPC system to analyze all videos using SLEAP’s command-line interface. The analysis was conducted on NVIDIA A100 GPUs with the following configuration: two nodes, three GPUs per node, and 128 GB of memory per GPU.

The SLURM batch scripts used for HPC analysis are provided in the NeuroPupil GitHub repository (https://github.com/KemalOz21/NeuroPupil) for reference and transparency. These scripts reflect the job submission workflow and computing environment used at the Cleveland Clinic Research. Because HPC infrastructures, scheduling systems, and resource management policies vary across institutions, HPC analysis workflows are not standardized. As such, the provided scripts are intended as reference implementations and may require modification to accommodate different institutional HPC systems, including adjustments to resource allocation, module loading, and environment configuration. The inclusion of these scripts is intended to illustrate how NeuroPupil analyses can be efficiently scaled in an HPC environment and to assist users in adapting the pipeline to their own computational resources.

### Facemap vs NeuroPupil Mouse

Facemap is a recently developed code package primarily used to analyze mouse facial video data^7^. In addition to facial analysis, Facemap’s GUI includes a module for automatic pupil detection and area measurement using an image augmentation strategy similar to that of Cellpose 2.0^34^. To evaluate the accuracy of pupil area estimation, the performance of the Animal NeuroPupil model was compared to Facemap across clean and challenging datasets from both the Wheel and Airball conditions (total of 40 videos, 20 from each experiment). Facemap version 1.0.7 was used to compute the Pupil areas from its automated detection and from the Animal NeuroPupil model by calculating the average diameter between the Top-Bottom and Right-Left labeled coordinates and subsequently estimating the area of the corresponding circle (**Fig. 2A-D**).

For ground-truth validation, four independent annotators manually measured pupil areas in 10 evenly sampled frames per video using ImageJ. A custom MATLAB script was then used to calculate the percentage error in area estimation for both the Animal NeuroPupil model and Facemap, with results plotted for each animal (**Fig. 2E**) and aggregated across all experiments (**Fig. 2F**). Runtimes for each experiment (**Fig. 2G**) were recorded for NeuroPupil on the local computer (**Supp. Fig. 13A**) and for Facemap. For the total runtime across all experiments (**Fig. 2H**), results from these two methods were plotted alongside NeuroPupil utilizing the high-performance computer (**Supp. Fig. 13B**) to access processing efficiency.

### Public and CCF Human Data Analysis

Two separate human pupil tracking datasets were analyzed using global SLEAP models following procedures similar to those developed for the animal datasets.

The first dataset (**Fig. 3A and Supp. Fig. 14A**), referred to as the public human dataset, was obtained from a previously published study^24^ and consisted of 22 patients, each undergoing three sessions, resulting in a total of 66 videos. For model construction, 500 frames were evenly extracted from each video and concatenated into a single combined video using the **mediaGUI** (**Supp. Fig. 19**). Two frames per video were labeled in SLEAP, using the same methodology established in the TMC animal models, to train a global model capable of analyzing all videos within the public human dataset. Analysis and labeled video generation were performed following the same procedures used for the animal datasets, with precision values for pupil area estimation plotted as averages across each patient (**Fig. 3B, top**) and for each individual experiment (**Supp. Fig. 15**).

The second dataset, referred to as the Cleveland Clinic Foundation (CCF) dataset (**Fig. 3A and Supp. Fig. 14B**), was collected from the Cleveland Clinic Cole Eye Institute and included 33 patients who underwent varying numbers of sessions, totaling 590 videos. For this dataset, 50 frames were evenly extracted from each video to construct a concatenated training video using the **mediaGUI** (**Supp. Fig. 19**). Similarly, two frames per video were labeled in SLEAP to create a global model capable of analyzing the entire CCF human dataset. Subsequent analysis and visualization followed the same workflow used for both the animal datasets and the public human dataset, with precision values for pupil area estimation plotted as averages across each patient (**Fig. 3B**) and for each individual experiment (**Supp. Fig. 16**). The runtimes were recorded for both model analysis (**Fig. 3E**) to access processing efficiency.

### NeuroPupil Human Analysis

The NeuroPupil Human model (**Fig. 3C and Supp. Fig. 14C**) was developed by combining concatenated videos from both the public and CCF human datasets. For model training, two frames per video were labeled, resulting in a dataset where most frames (90%) originated from the CCF dataset due to its substantially larger number of videos (n = 590) compared to the public dataset (n = 66). The model was trained in SLEAP using the same optimized parameters we established for the previous mouse SLEAP models. Following training, NeuroPupil Human maintains high precision on CCF data (which comprises 90% of training) and, importantly, generalizes well to the public dataset despite limited exposure during training (10% of training data) (**Fig. 3D**, **Supp. Fig. 17** for the **public dataset and Supp. Fig. 18** for the **CCF dataset**).

To evaluate processing efficiency, runtimes for the Human NeuroPupil model were recorded (**Fig. 3E**) utilizing the NeuroPupil local computer (**Supp. Fig. 19A**) and the high-performance computer analysis (**Supp. Fig. 19B**) method environment. Timing measurements included the complete workflow: video concatenation, frame labeling, model training, and video analysis for both the public and CCF datasets.

### Facemap vs NeuroPupil Human

A similar procedure that was done to compare the Facemap code package to NeuroPupil Animal was done for the Human NeuroPupil model. Clean and challenging datasets from both the Public and CCF human conditions (total of 14 videos; 6 from Public and 8 from CCF) were analyzed (**Fig. 4A-D**). For ground-truth validation, four independent annotators manually measured pupil areas in 10 evenly sampled frames per video using ImageJ. A custom MATLAB script was then used to calculate the percentage error in area estimation for both the Human NeuroPupil model and Facemap, with results plotted for each patient (**Fig. 4E**) and aggregated across all experiments (**Fig. 4F**). Runtimes for each experiment (**Fig. 4G**) were recorded for NeuroPupil on the local computer (**Supp. Fig. 19A**) and for Facemap. For the total runtime across all experiments (**Fig. 4H**), results from these two methods were plotted alongside NeuroPupil utilizing the high-performance computer (**Supp. Fig. 19B**) to access processing efficiency.

### Brain Activity Prediction Model from Pupil Keypoint Features

Five GCaMP6s-expressing mice underwent six days of behavioral and neural imaging, yielding a total of 30 paired pupil–brain datasets. Each pupil video was recorded at 40 Hz and accompanied by GCaMP6s fluorescence sampled at 20 Hz across ten brain regions in both hemispheres. Two frames per pupil video were manually annotated at four keypoints (Top, Bottom, Left, Right), and these identical annotations were used to train both NeuroPupil and DeepLabCut (DLC) for consistent pupil tracking across methods. Facemap was processed using its built-in unsupervised pipeline. For all models, pupil features sampled at 40 Hz were resampled onto the 20 Hz GCaMP time base using linear interpolation. Any remaining missing samples (such as dropped frames) were then filled using shape-preserving piecewise cubic Hermite interpolation (pchip). Each brain signal was smoothened with a 5 s moving mean filter prior to modeling^35^.

NeuroPupil and DLC each produced four pupil keypoints with x and y coordinates, yielding an eight-dimensional feature vector (x,y coordinates for the Top (𝑥_𝑇_, 𝑦_𝑇_), Bottom (𝑥_𝐵_, 𝑦_𝐵_) Right (𝑥_𝑅_, 𝑦_𝑅_) and Left (𝑥_𝐿_, 𝑦_𝐿_)) at each timepoint t:

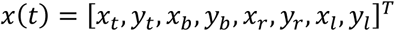

Preprocessing was performed after insertion of the eight raw coordinate parameters and included removal of score/index columns (NeuroPupil), correction of missing coordinates using pchip interpolation, and extraction of geometric descriptors including the x- and y-centroid, diameter and its temporal derivative, which were smoothed using a 3 second moving average at 40Hz to reduce noise. Afterwards, the 40 Hz pupil was resampled to 20 Hz to match the sampling rate of the brain signal prior to model training. A feedforward neural network which was trained to predict brain activity using the nonlinear mapping is,

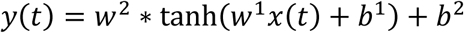

where 𝑥(𝑡) is the input, 𝑤^1^, 𝑤^2^ are the weights, 𝑏^1^, 𝑏^2^ are the biases of the model and as an activation function we choose tanh(·). To optimize this model, we used Levenberg–Marquardt backpropagation to minimize Mean Squared Error (MSE). A total of 27 experiment pairs were used for training and 3 for testing. Performance was quantified using Root Mean Squared Error (RMSE), Coeficient of Determination (R²), and 1% trimmed RMSE/R² (removing the top 1% of absolute errors). Architecture sweeps were conducted using 5, 10, 15, and 20 hidden neurons with 50–2000 epochs. The best-performing NeuroPupil model used 15 hidden neurons and 100 epochs, while DLC performed best with 5 hidden neurons and 250 epochs. With eight input features, the motor (MOp), somatosensory (SSp_ll, SSp_ul, SSp_tr), retrosplenial (RSPagl, RSPd, RSPv) and visual (VISa,VISam, VISpm) regions were successfully predicted. A two-sample t-test was applied to compare region-wise R² values between NeuroPupil and DLC.

The same preprocessing, temporal alignment, smoothing, training/testing split, neural network architecture, and performance metrics were applied to the NeuroPupil-vs-Facemap comparison. The only methodological difference was the construction of the input feature vector. Facemap outputs a single pupil-area value; therefore, NeuroPupil’s four keypoints were reduced to an equivalent scalar by computing the ellipse area *A(t)* from the major *a(t)* and minor *b(t)* axes. Let

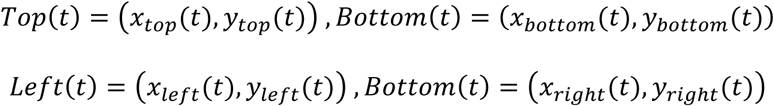

denote the tracked pupil boundary coordinates at time *t*. The major and minor axis were computed as half of the Euclidan distances between opposing boundary points:

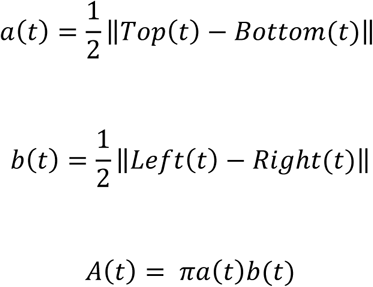

This produced a one-dimensional input vector,

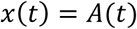

which was inserted into the same feedforward network architecture used for the eight-parameter coordinate model, ensuring comparable model structure despite the reduced feature dimensionality.

Architecture sweeps were identical, and performance again was evaluated with RMSE, R², and 1% trimmed metrics. The optimal NeuroPupil-area model used 10 hidden neurons and 500 epochs, while Facemap performed best with 10 hidden neurons and 300 epochs. In contrast to the eight-feature model, the one-parameter model successfully predicted only six of the ten brain regions (MOp, RSPagl, RSPv1, SSp_ll, SSp_ul, SSp_tr, and VISam). Region-wise R² values for NeuroPupil-area and Facemap were compared using a two-sample t-test.

Across both comparisons, these studies demonstrate that increased input dimensionality dramatically improves brain-activity prediction performance. The eight-parameter NeuroPupil model preserved richer geometric and spatial information about pupil behavior and successfully predicted all ten brain regions. In contrast, reducing the model to a single area feature, as required for comparisons with Facemap, substantially limited predictive capacity and resulted in accurate predictions in only six regions. Therefore, multimodal or higher-dimensional representations of pupil dynamics provide a more powerful and nuanced description of brain state than single-parameter models.

### Patient-level disease classification using a unified 1D convolutional neural network

#### Data organization and signal sources

Pupil signals were extracted from the same subject cohort using three independent processing pipelines: NeuroPupil, DeepLabCut (DLC), and Facemap. Subjects were categorized into Healthy and Disease groups, with recordings acquired on up to three experimental days per subject. NeuroPupil and DLC provided time-resolved pupil landmark trajectories, whereas Facemap yielded region-of-interest (ROI) pupil area time series. To enable a fair and method-independent comparison, all signals were transformed into a unified one-dimensional representation prior to analysis which is elliptic area.

#### Patch extraction

To obtain fixed-length inputs for learning, pupil area signals were segmented into non-overlapping temporal patches of 500 samples (𝐿 = 500). Patch extraction was performed deterministically. Patches containing non-finite or invalid values were excluded to ensure numerical stability and comparability across methods.

#### Patient-level cross-validation design

The dataset comprised 32 subjects, including 16 Healthy and 16 Disease individuals. Each subject contributed recordings from up to three experimental days. For each subject, pupil signals were segmented into non-overlapping patches of 500 samples, and exactly four patches were selected from each recording day. Consequently, each subject contributed a total of 12 patches (3 days × 4 patches/day). Subjects who did not meet this criterion due to insufficient valid data were excluded from the analysis. This deterministic sampling strategy ensured uniform contribution across subjects and controlled for inter-day variability. To avoid information leakage, all patches originating from a given subject were assigned exclusively to a single fold. Model evaluation was performed using balanced patient-level 8-fold cross-validation. In each fold, the test set consisted of 4 subjects, 2 Healthy and 2 Disease, while the remaining 28 subjects (14 Healthy, 14 Disease) were used for training and validation. Across the eight folds, each subject served as a test subject exactly once, yielding a complete and unbiased assessment of generalization performance at the patient level.

#### <u>Model architecture</u>: 1D convolutional neural network

Patch-level classification was performed

using a lightweight one-dimensional convolutional neural network (1D-CNN). Each input patch was a single-channel normalized pupil signal 𝑥 ∈ ℝ^500×1^. To increase representational capacity prior to convolution, the input was first projected at each time point from one feature to 32 channels using a linear layer applied along the feature dimension.

The network consisted of three sequential strided convolutional blocks:

1. **Block 1:** Conv1D (32 → 32) with kernel size 9, stride 2, and padding 4, followed by batch normalization, ReLU activation, and dropout (𝑝 = 0.2).
2. **Block 2:** Conv1D (32 → 64) with kernel size 7, stride 2, and padding 3, followed by batch normalization, ReLU activation, and dropout (𝑝 = 0.2).
3. **Block 3:** Conv1D (64 → 128) with kernel size 5, stride 2, and padding 2, followed by batch normalization and ReLU activation.

Strided convolutions progressively reduced the temporal resolution (500 → 250 → 125 → 63) while increasing channel capacity. Following the final convolutional block, global average pooling was applied over the temporal dimension to produce a fixed-dimensional feature vector in ℝ^128^. A final linear layer mapped this representation to a single output logit for binary classification (Healthy vs Disease).

#### Model training and optimization

Models were trained using the AdamW optimizer with a learning rate of 10^−3^ and weight decay of 10^−4^. To address class imbalance at the patch level, binary cross-entropy loss with logits was employed with class weighting based on the training data distribution. Learning rate reduction on plateau and early stopping based on validation loss were applied to prevent overfitting. Gradient norm clipping was used to enhance training stability.

#### Evaluation metrics

Model performance was evaluated on held-out test subjects using area under the ROC curve (AUC), accuracy, precision, recall, specificity, and F1 score. Performance metrics were computed at the patch level and aggregated across folds.

## SUPPLEMENTARY FIGURES

**Supplementary Figure 1.**
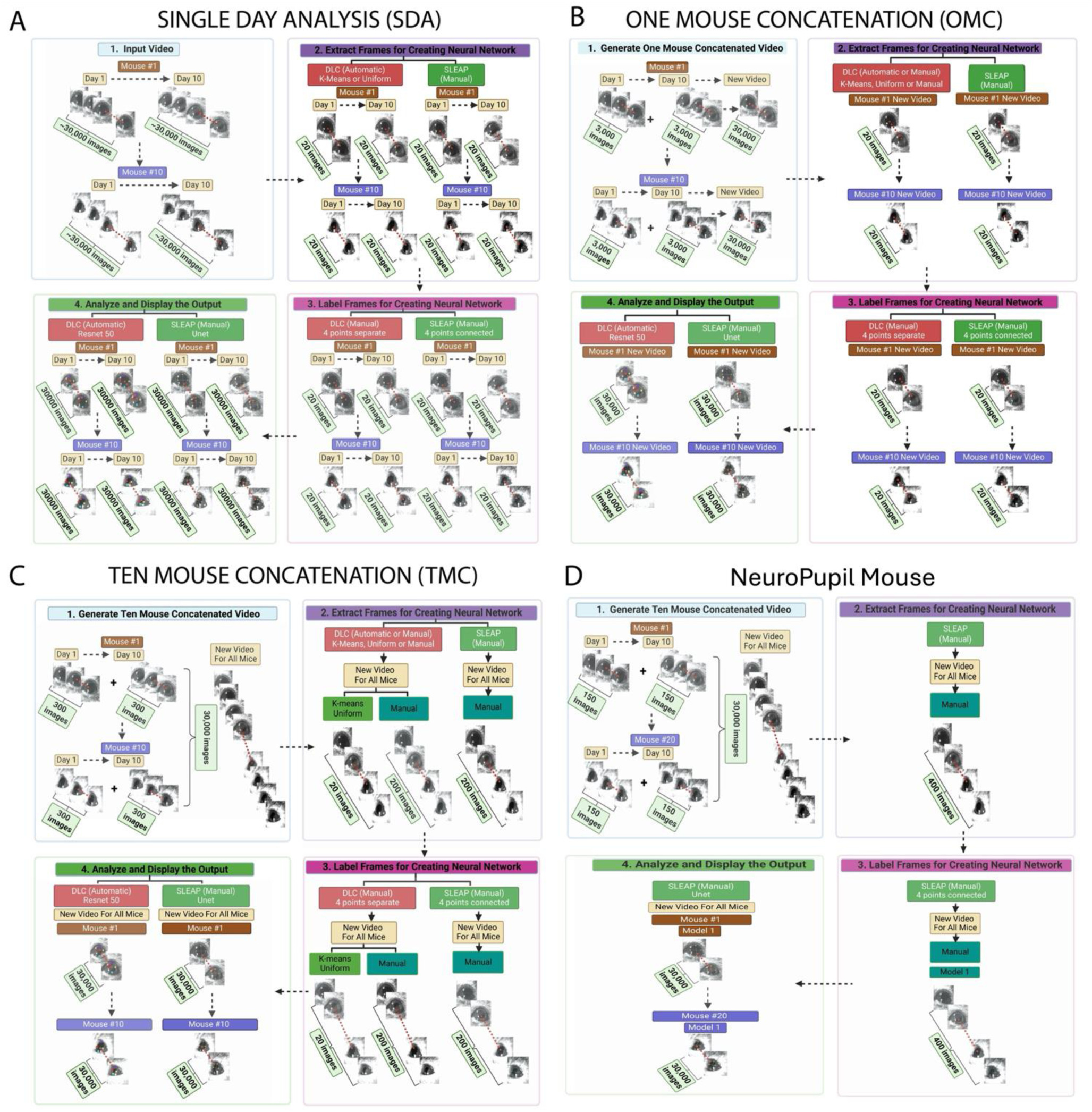
Detailed Workflow of Model Training Methods for the Mouse Experimental Data. **(A) Single Day Analysis (SDA). Workflow illustrating model generation using the SDA approach**. First, (1) an experimental video was loaded into the code package (DLC or SLEAP). (2) Frames were then extracted (n = 20) to create the model. DLC applied unsupervised frame extraction (k-means or uniform), while SLEAP used evenly spaced manual extraction. (3) Extracted frames were labeled to train the machine learning model (ResNet50 for DLC, U-Net for SLEAP), with four pupil landmarks (Top, Bottom, Right, Left) annotated in each frame. In the SLEAP package, labeled points could also be connected to help the model better learn spatial relationships between keypoints; therefore, the Top point was connected to the Bottom point, and the Right point was connected to the Left point. (4) Once trained, the model analyzed the input video to predict pupil positions across all frames. Outputs included a CSV file with predicted (x, y) coordinates and confidence scores for each keypoint, as well as a labeled video visualizing prediction accuracy**. (B) One Mouse Concatenation (OMC)**. Workflow illustrating model generation using the OMC approach. To accelerate video analysis, a model was created per animal by (1) extracting 3,000 frames from each experimental video (n = 10 per animal) and concatenating them into a 30,000-frame composite video. (2) This concatenated video was processed in the same code packages as in SDA, with frame extraction performed identically. To match SLEAP’s two-frames-per-video protocol, an additional evenly spaced extraction method was applied in DLC. (3) Extracted frames were labeled and models were trained as before. (4) These models were then applied back to the original experimental videos (used to generate the concatenated dataset), producing the same output files as in the SDA workflow**. (C) Ten Mouse Concatenation (TMC)**. Workflow illustrating model generation using the TMC approach. To further accelerate video analysis, a model was created using all experimental videos (wheel and airball) by (1) extracting 300 frames from each video (n = 100 per experiment) and concatenating them into a 30,000-frame composite video. (2–3) The concatenated video was then processed similarly to the OMC workflow. For DLC, 20 frames were labeled using k-means or uniform extraction, while for manual models in DLC and SLEAP, 200 images were labeled to ensure two frames were annotated per experimental video. (4) The trained models were then applied to the original experimental videos used to generate the concatenated dataset. **(D) NeuroPupil Mouse Model. Workflow illustrating the development of NeuroPupil Mouse**. After SLEAP was identified as the most robust platform for pupil analysis across both experimental conditions, a generalized model was created to analyze pupillometry data not only from wheel and airball experiments, but also from different animal types (e.g., PTEN, MS, and GBM). To build the model, (1) 150 frames were extracted from each video using mediaGUI (Supp. Fig. 2) and combined into a concatenated dataset spanning wheel (n = 100) and airball (n = 100) experiments. (2–3) This dataset was imported into SLEAP, where two images per day (n = 400) were labeled for training. Once trained, NeuroPupil was used to analyze wheel and airball data either via the custom-built sleapGUI (**Supp.** Fig. 13A) or on the HPC system (**Supp.** Fig. 13B).

**Supplementary Figure 2.**
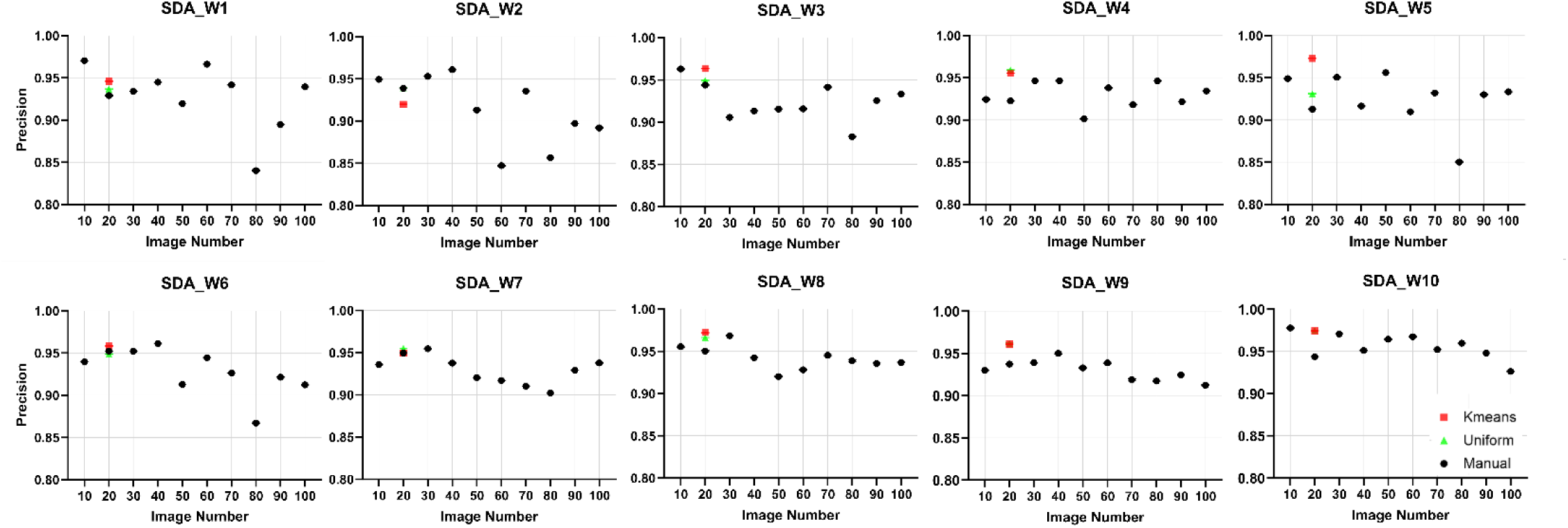
DeepLabCut SDA Image Extraction. For each animal in the low-resolution wheel dataset, one experimental video was selected, and SDA models were generated using image extraction ranges from 10 to 100 frames. Manually extracted images were obtained in an evenly spaced manner. By default, DLC’s k-means and uniform unsupervised image extraction algorithms extract 20 frames. The results showed that high precision values were achieved, and these did not change significantly as the number of labeled images increased. Therefore, k-means and uniform extraction methods were used to perform SDA analysis for both experimental conditions.

**Supplementary Figure 3.**
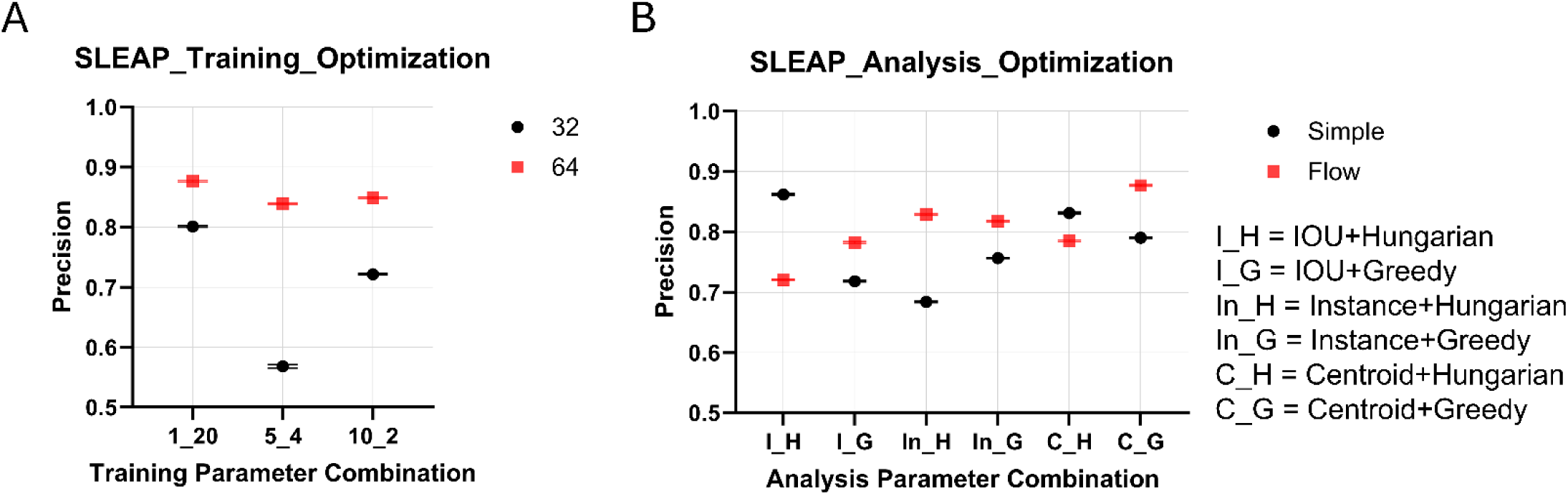
SLEAP Training and Analysis Optimization. **(A) Training Parameter Optimization**. SLEAP v1.3.3 was tested on a concatenated video from subject F11 (wheel dataset, Day 10) to determine optimal parameters. Three K-means clustering settings (1, 5, 10 clusters; 20 frames total so we have 1_20:1 cluster and 20 frames each, 5_4: 5 clusters and 4 frames each, 10_2: 10 clusters and 2 frames each) and two frame sizes (32, 64) were evaluated using the U-Net architecture with default post-processing. The highest precision was achieved with 20 frames from a single cluster and frame size 64. Based on these results, subsequent models used even frame selection rather than clustered sampling**. (B) Analysis Parameter Optimization**. After determining optimal training settings, the same video used to optimize the training parameters was used to evaluate analysis parameters. Two tracking methods (Simple, Flow), three instance matching strategies (IOU, Instance, Centroid), and two assignment algorithms (Greedy, Hungarian) were systematically tested, resulting in 12 parameter combinations. The highest precision combination was obtained with Flow tracking, Centroid matching, and Greedy assignment.

**Supplementary Figure 4.**
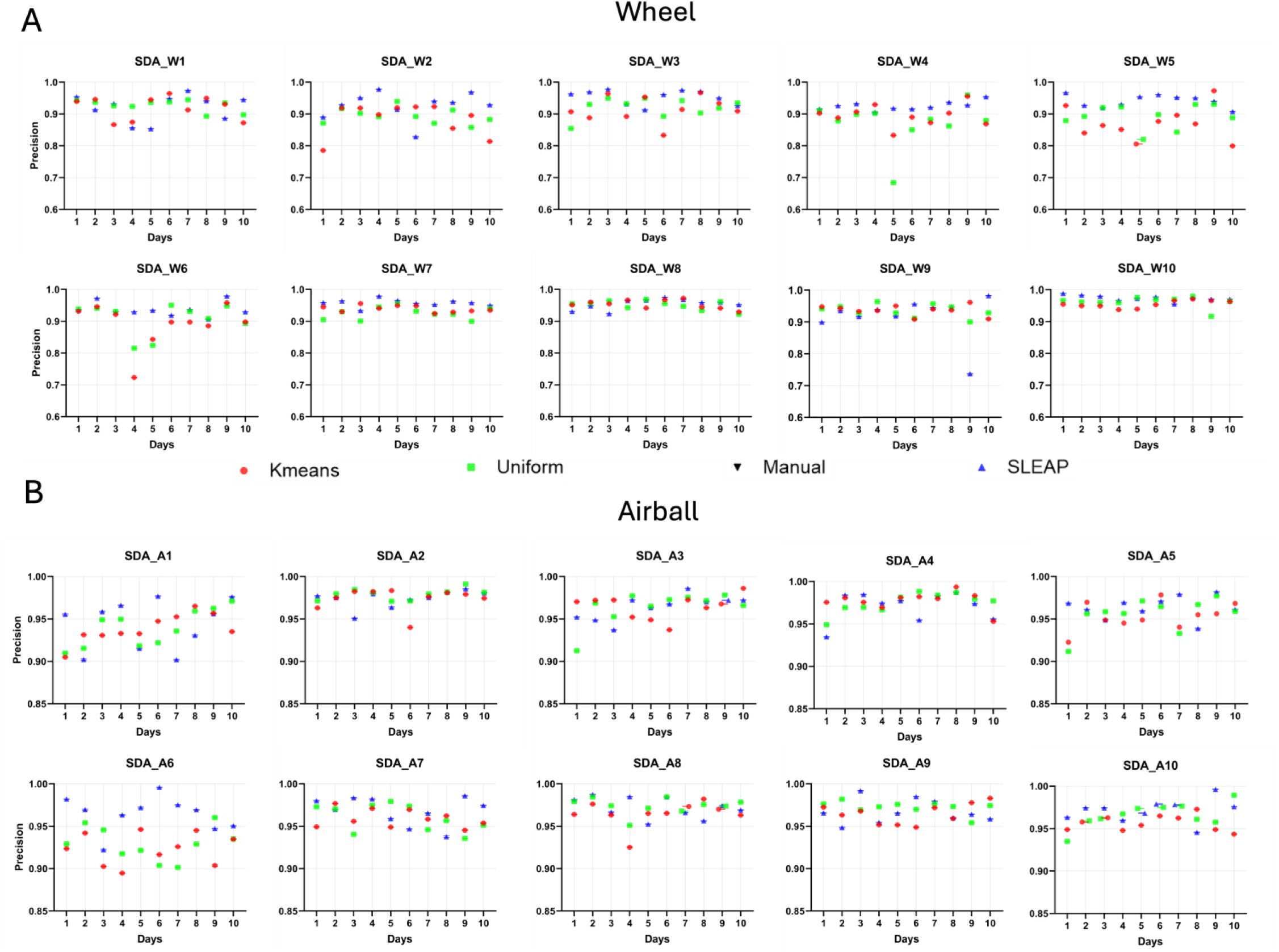
Detailed SDA Precision Values. **(A) SDA on the Wheel Experiment**. Detailed precision values were done for the low-resolution wheel experiment using the SDA method. In each mouse (n = 10) across 10 experimental days, the precision and standard error were plotted on each day for the y axis with respect to each model. DeepLabCut (kmeans and uniform) and SLEAP. **The same SDA precision analysis was performed for the (B) high-resolution Airball Experiment**, which included data from 10 animals across 10 different experimental days.

**Supplementary Figure 5.**
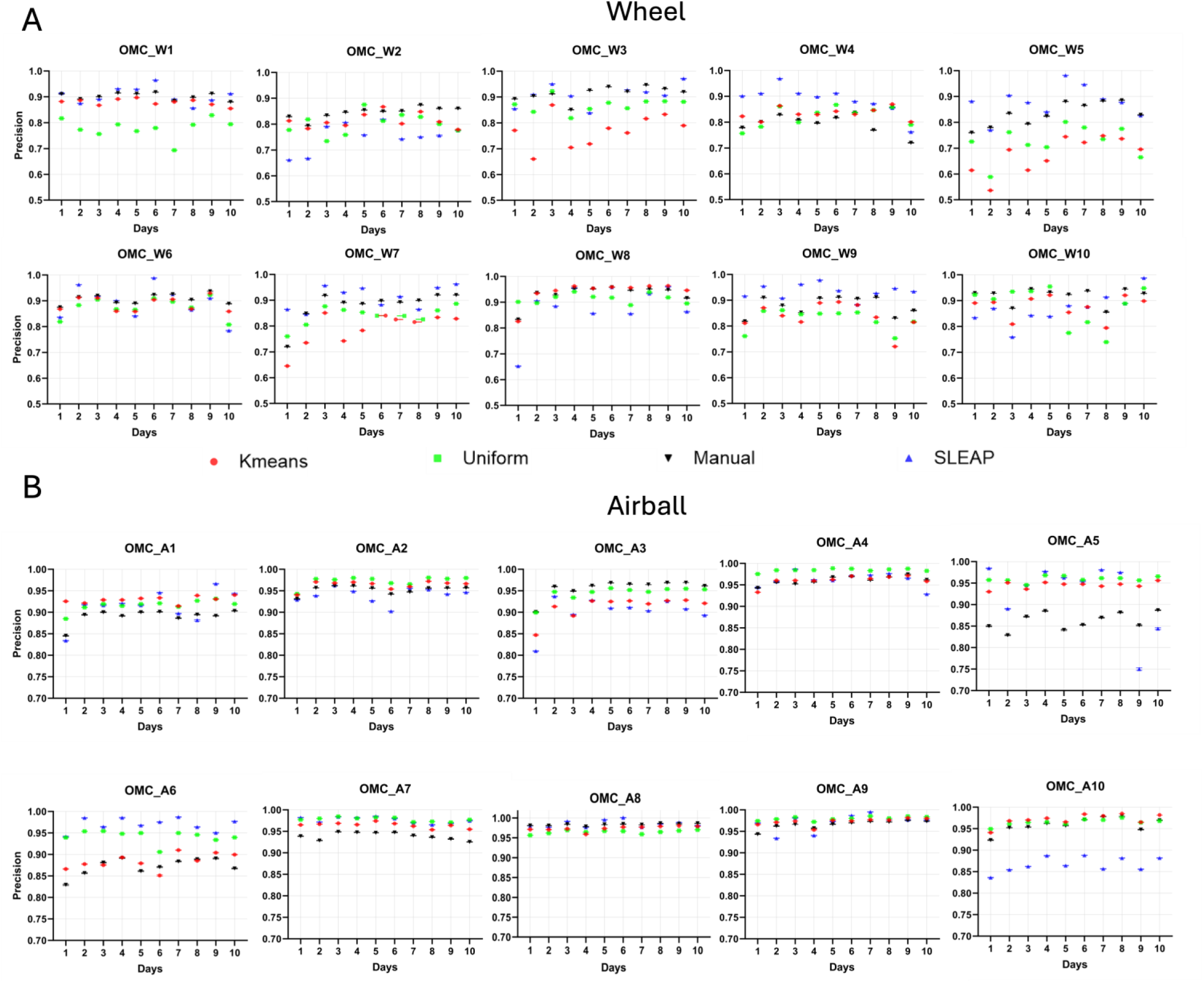
Detailed OMC Precision Values. **(A) OMC Analysis on the Wheel Experiment**. Detailed precision values were done for the low-resolution wheel experiment using the OMC method. In each mouse (n = 10) across 10 experimental days, the precision and standard error were plotted on each day for the y axis with respect to each model: DeepLabCut (kmeans, uniform, manual) and SLEAP. **The same OMC precision analysis was performed for the (B) high-resolution Airball Experiment**, which included data from 10 animals across 10 different experimental days.

**Supplementary Figure 6.**
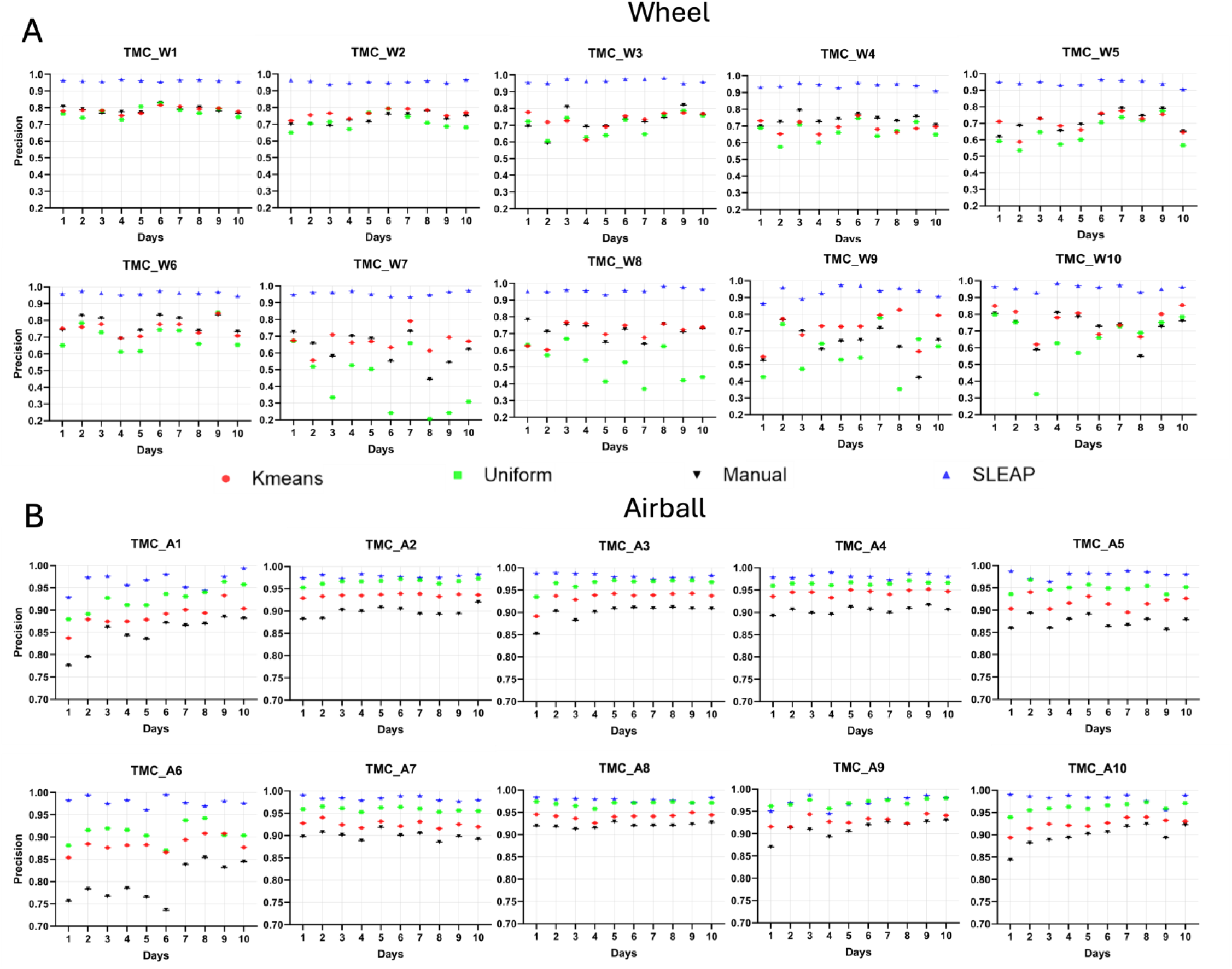
Detailed TMC Precision Values. (A) TMC Analysis on the Wheel Experiment. Detailed precision values were done for the low-resolution wheel experiment using the TMC method. In each mouse (n = 10) across 10 experimental days, the precision and standard error were plotted on each day for the y axis with respect to each model: DeepLabCut (kmeans, uniform, manual) and SLEAP. **The same SDA precision analysis was performed for the (B) high-resolution Airball Experiment,** which included data from 10 animals across 10 different experimental days.

**Supplementary Figure 7.**
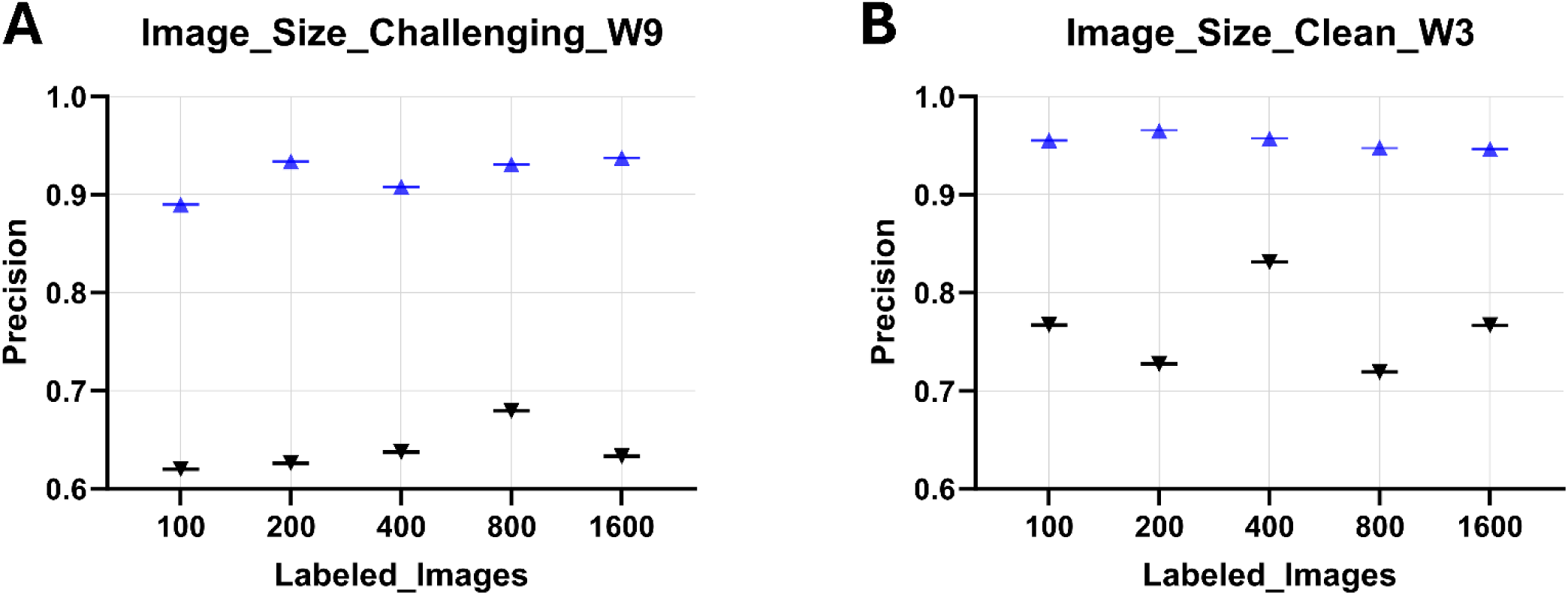
Effect of number of labeled images on TMC model precision in SLEAP (blue) and DLC (black). Precision was evaluated for Ten-Mouse Concatenation (TMC) models trained with increasing numbers of labeled images (100, 200, 400, 800, and 1,600 total images)**. (A)** Challenging video (W9) and **(B)** clean video (W3) from the wheel dataset. Blue markers indicate NeuroPupil (SLEAP, U-Net) performance, while black markers indicate DeepLabCut (ResNet-50) performance. Precision gains plateaued after approximately 200 labeled images (corresponding to 2 images per video), with minimal improvement beyond this point. The wheel dataset was used for this analysis because its lower spatial resolution makes it more challenging than the airball experiments.

**Supplementary Figure 8.**
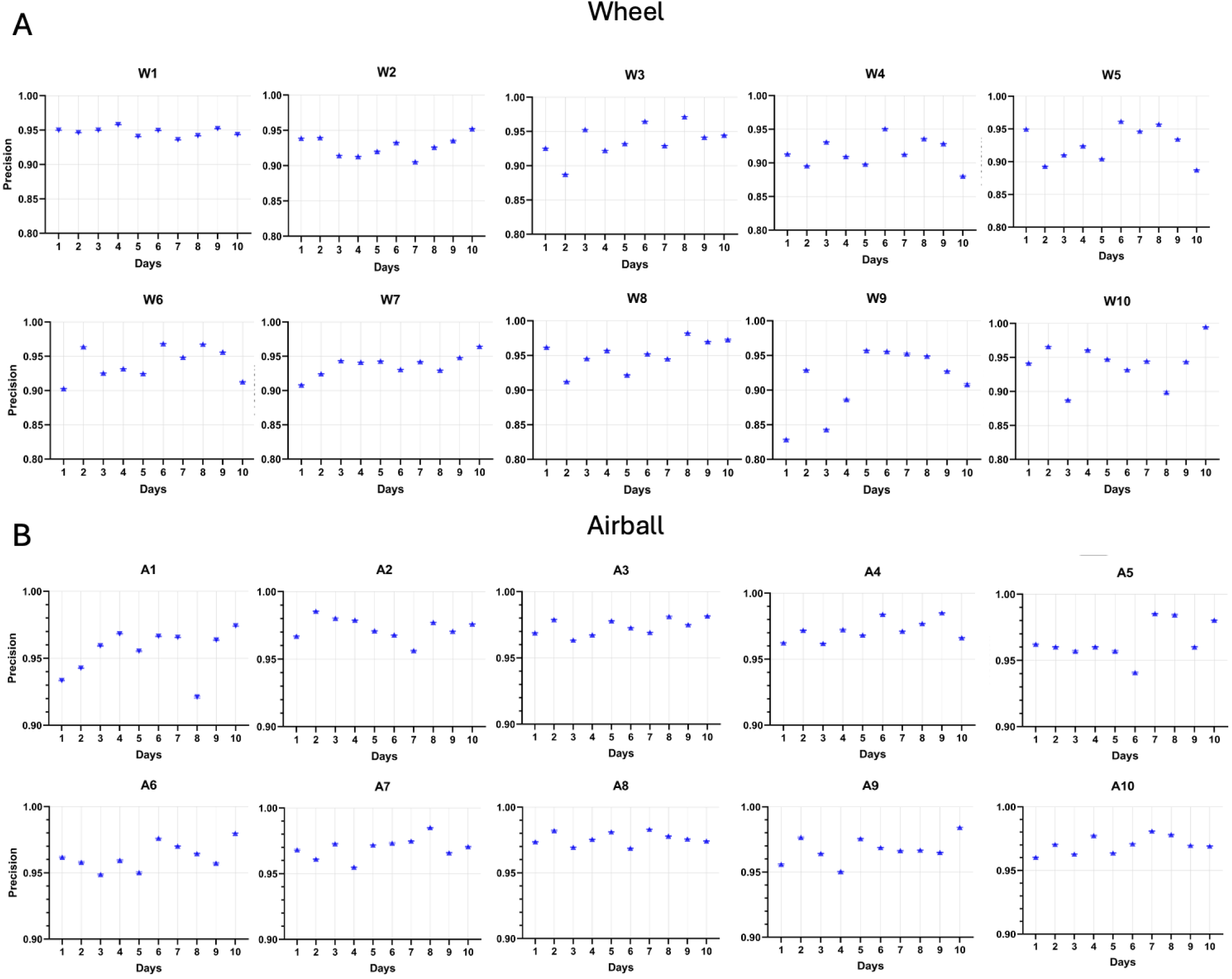
Detailed NeuroPupil Precision Values for Wheel and Airball Data. **(A) NeuroPupil Analysis on the Wheel Experiment**. Detailed precision values were done for the low-resolution wheel experiment using the NeuroPupil method. In each mouse (n = 10) across 10 experimental days, the precision and standard error were plotted on each day for the y axis with respect the NeuroPupil model. **The same SDA precision analysis was performed for the (B) high-resolution Airball Experiment**, which included data from 10 animals across 10 different experimental days.

**Supplementary Figure 9.**
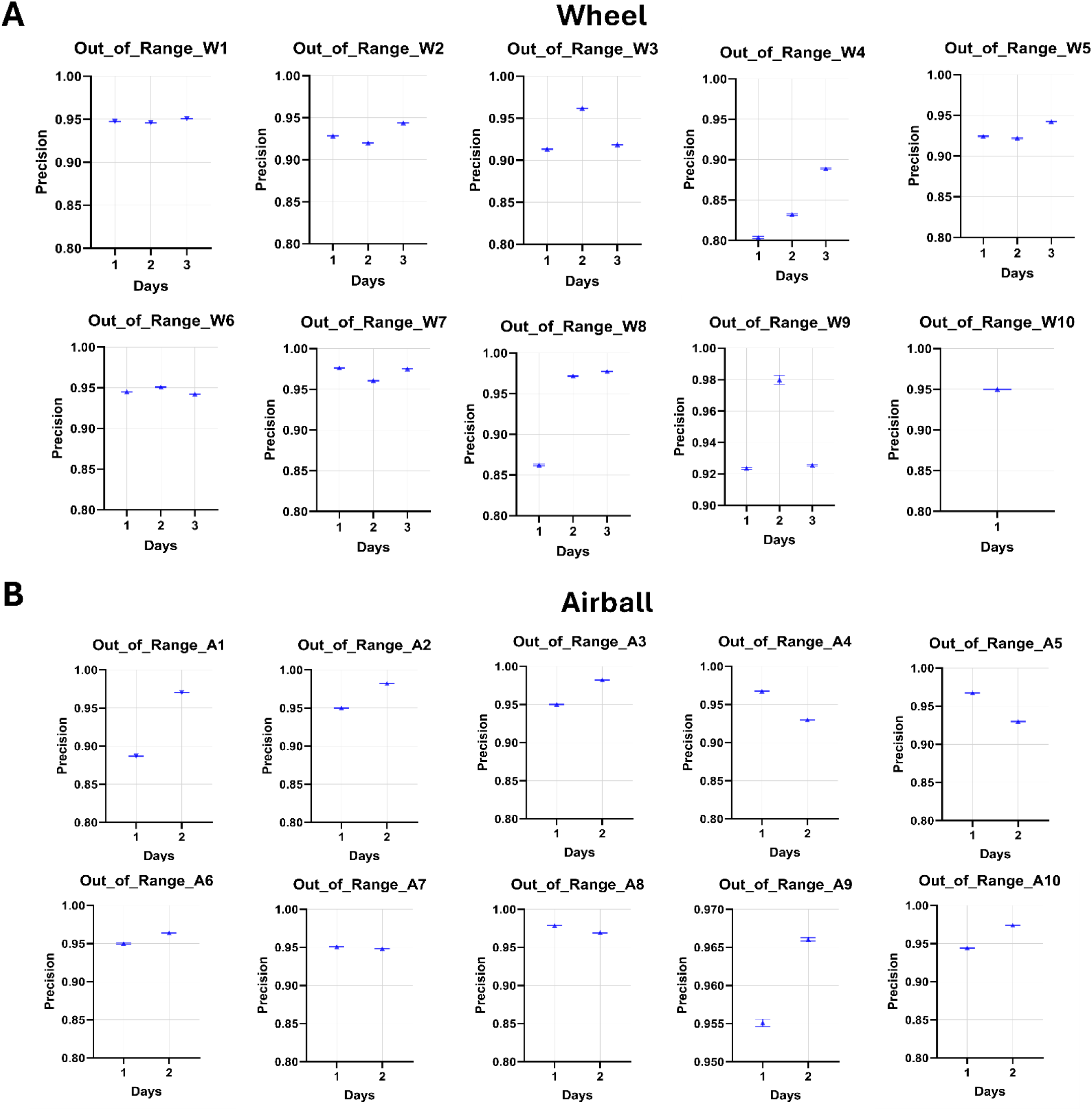
NeuroPupil Precision Values for Out-of-Range Wheel and Airball Data. **(A) Wheel Experiment**. Precision values were evaluated for the low-resolution wheel dataset, which was not included in NeuroPupil model training. For each mouse (n = 10) across multiple experimental days, precision and standard error were plotted by day relative to the NeuroPupil model**. (B**) **Airball Experiment.** The same SDA precision analysis was performed for the high-resolution airball dataset, which also included data from 10 animals across several experimental days that were not part of the NeuroPupil training set.

**Supplementary Figure 10.**
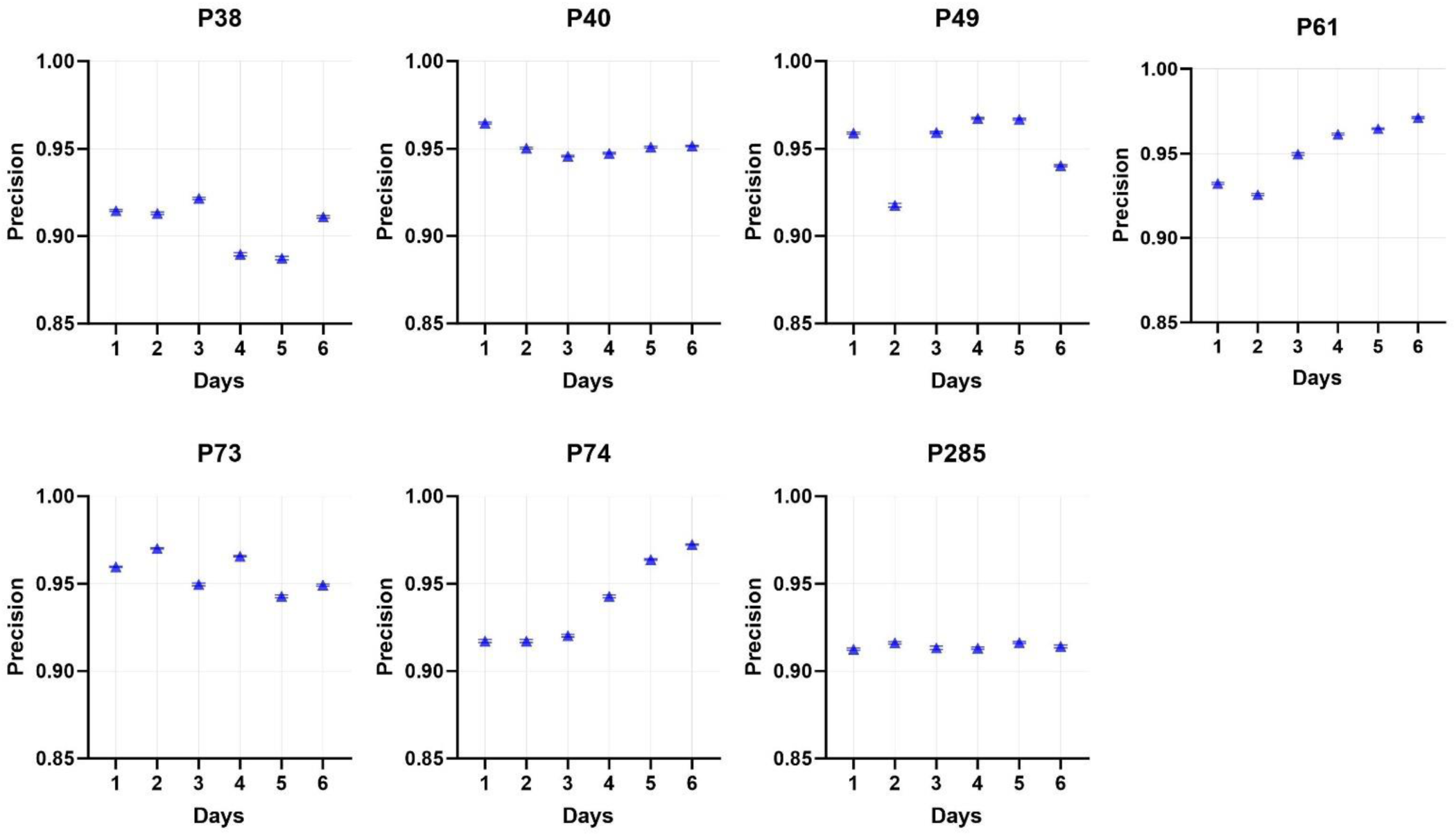
NeuroPupil Precision Values for PTEN Data. Precision values were evaluated for the PTEN dataset, which was not included in NeuroPupil model training. For each mouse (n = 7) across multiple experimental days, precision and standard error were plotted by day relative to the NeuroPupil model.

**Supplementary Figure 11.**
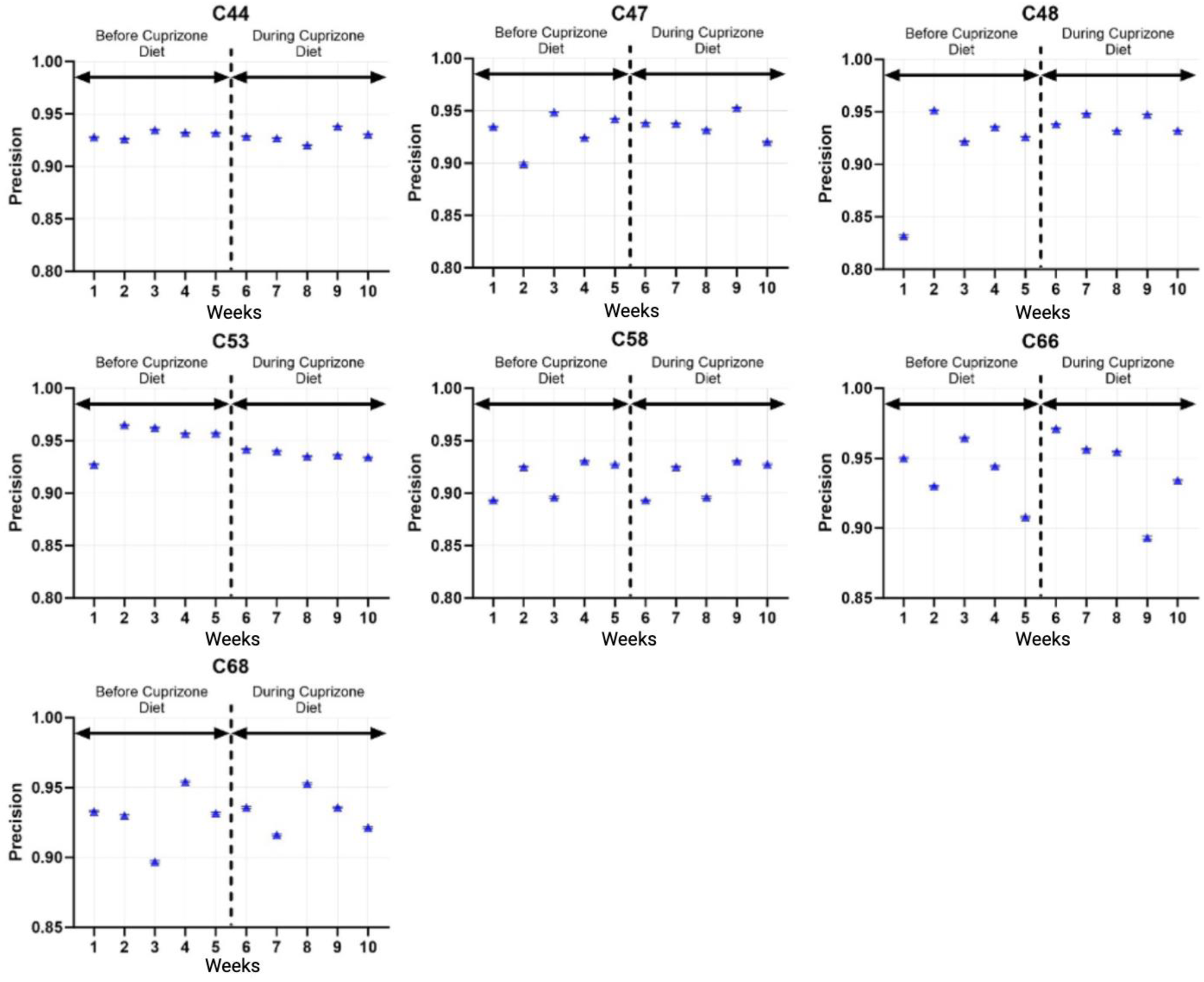
NeuroPupil Precision Values for Healthy and Cuprizone MS Data. Precision values were evaluated for the MS dataset, which was not included in NeuroPupil model training. For each mouse (n = 7) across 10 experimental days, precision and standard error were plotted by day relative to the NeuroPupil model. The first 5 days corresponded to a normal diet, while the remaining 5 days included a cuprizone diet.

**Supplementary Figure 12.**
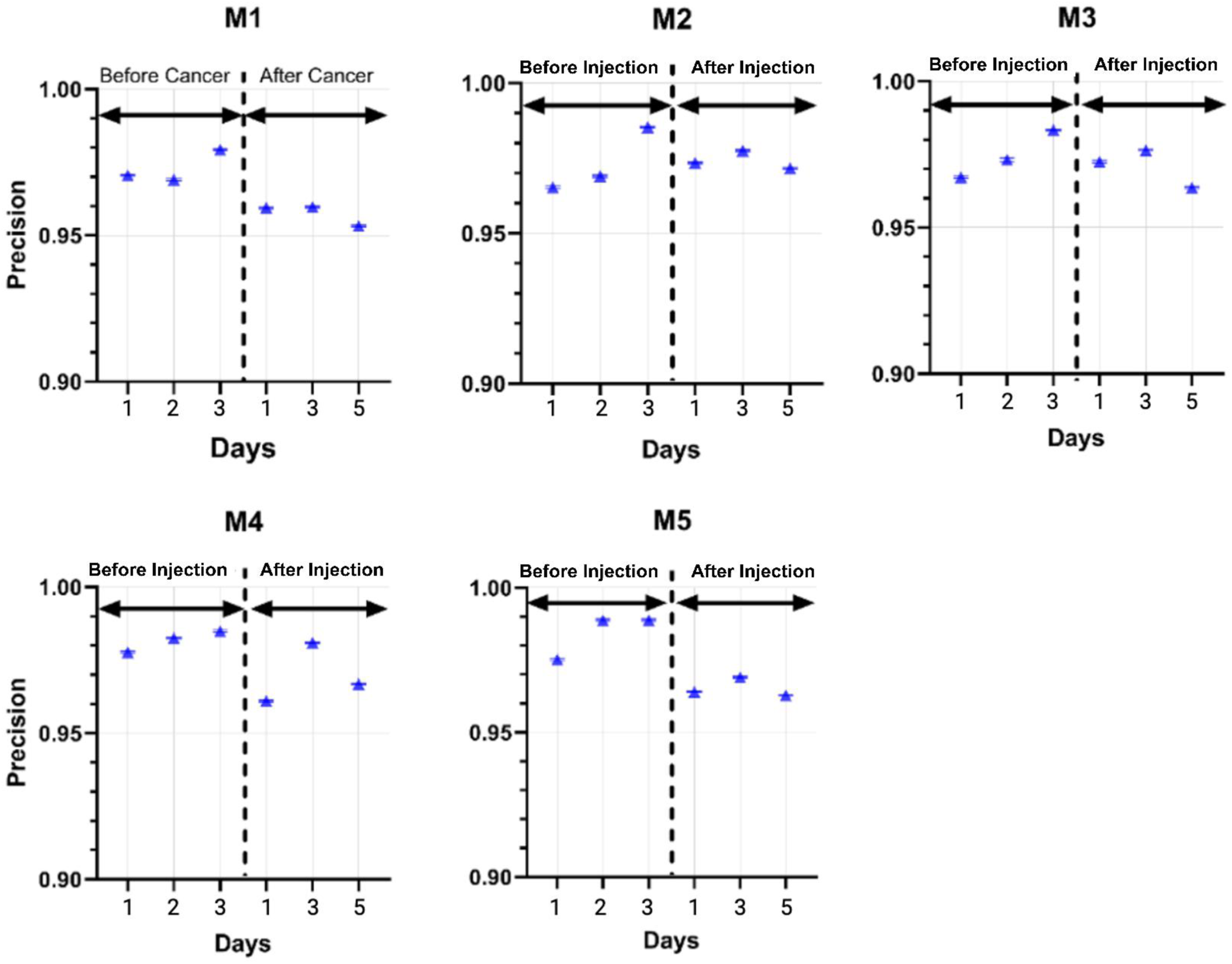
NeuroPupil Precision Values for before and after SB28 tumor injection for glioblastoma (GBM) modeling. Precision values were evaluated for the GBM dataset, which was not included in NeuroPupil model training. For each mouse (n = 5 in total) across 6 experimental days, precision and standard error were plotted by day relative to the NeuroPupil model. The first 3 days corresponded to the pre-injection (healthy) condition, while the remaining 3 days followed after tumor (SB28) injection for glioblastoma modeling.

**Supplementary Figure 13.**
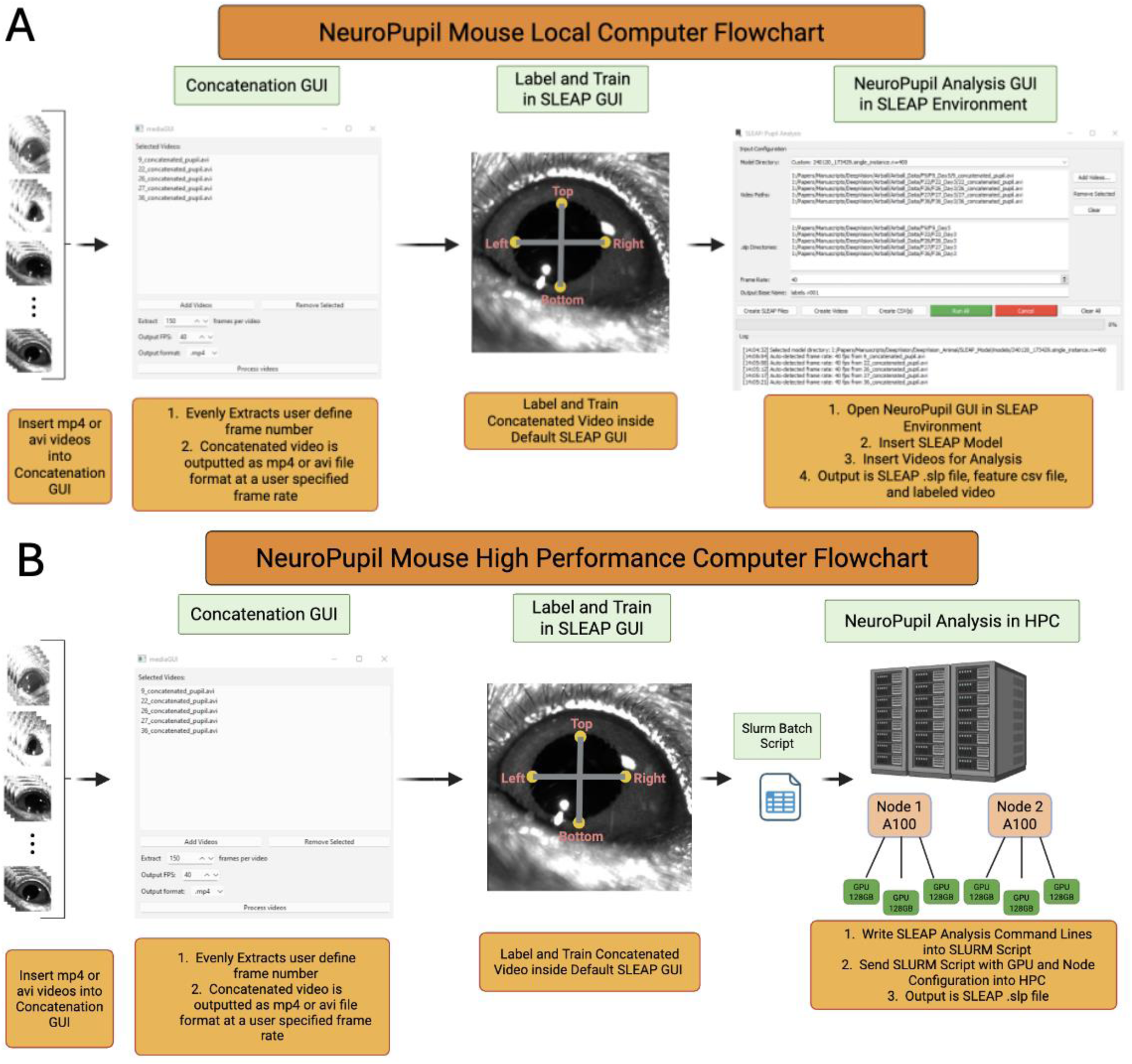
NeuroPupil Mouse Workflow for Local and High-Performance Computing Environments Caption. **(A) NeuroPupil Detailed Analysis Flow Chart for Local Computer**. All of the experimental pupil videos (either mp4 or avi video format) are inserted into the Concatenation GUI for creating a concatenated video (mp4 or avi format). The GUI will ask the user to specify the number of frames it will be extracted evenly per video, as well as the concatenated video frame rate and output location. After the video is concatenated, it is uploaded into the SLEAP default GUI for labeling and training the model. After training the model, the custom NeuroPupil Analysis GUI is opened inside the SLEAP Anaconda environment for analyzing the videos. Once the GUI opens, it requires the user to first input the SLEAP model location, then insert the desired videos into the video path location. After the videos are placed into the video path location, the GUI automatically generates the SLEAP analysis file (.slp), feature coordinates csv file, and labeled video inside the same file path as the inserted raw video. Finally, clicking the Run All green button will analyze all of the inserted video data**. (B) NeuroPupil Detailed Analysis Flow Chart for High Performance Computer**. The NeuroPupil analysis process is the exact same as in the local computer until after training the SLEAP model. Once training the model, a slurm batch script is created to analyze the video data. Inside the script, the number of nodes and GPUs are specified, as well as the SLEAP commands to analyze all of the videos. Then, the slurm batch script is sent to the HPC for video analysis and the output is the SLEAP analysis file

**Supplementary Figure 14.**
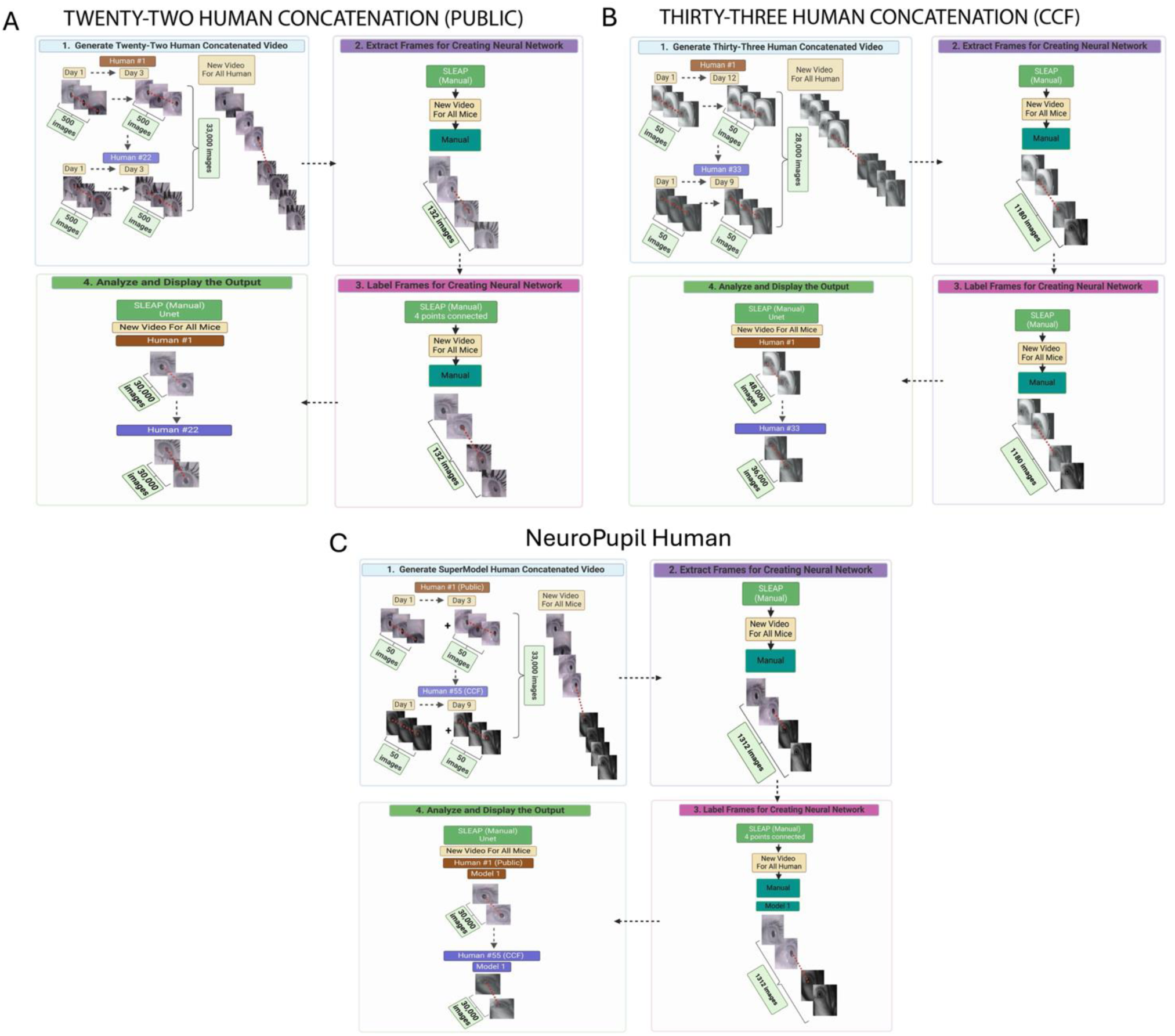
Detailed Workflow of Model Training Methods for the Human Experimental Data. (A) Public Human Model. Workflow illustrating model generation using the Public Human data. Similar to the TMC approach for the mouse data, a model was created using all experimental videos coming from the Public Human data by (1) extracting 500 frames from each video (n = 66) and concatenating them into a concatenated video. (2–3) The concatenated video was then processed similarly as the TMC method in the SLEAP code package, where two images were labeled per patient video. (4) The trained models were then applied to the original experimental videos used to generate the concatenated dataset. **(B) CCF Human Model. Workflow illustrating model generation using the CCF Human data**. Similar to the Public Model, a model was created using all experimental videos coming from the Cleveland Clinic patient data by (1) extracting 50 frames from each video (n = 590) and concatenating them into a concatenated video. (2–3) The concatenated video was then processed in the same manner as the Public Human model method in the SLEAP code package, where two images were labeled per patient video. (4) The trained models were then applied to the original experimental videos used to generate the concatenated dataset. **(C) NeuroPupil Human Model**. Workflow illustrating the development of NeuroPupil Human Model. A generalized model was created to analyze human pupillometry data coming from Public and CCF data. To build the model, (1) 50 frames were extracted from each video using mediaGUI (**Supp.** Fig. 19) and combined into a concatenated dataset for analyzing Public (n = 66) and CCF (n = 590) videos. (2–3) This dataset was imported into SLEAP, where two images per day (n = 1312) were labeled for training. Once trained, NeuroPupil was used to analyze the entire patient data via either the custom-built sleapGUI (**Supp.** Fig. 19a) or on the HPC system (**Supp.** Fig. 19b).

**Supplementary Figure 15.**
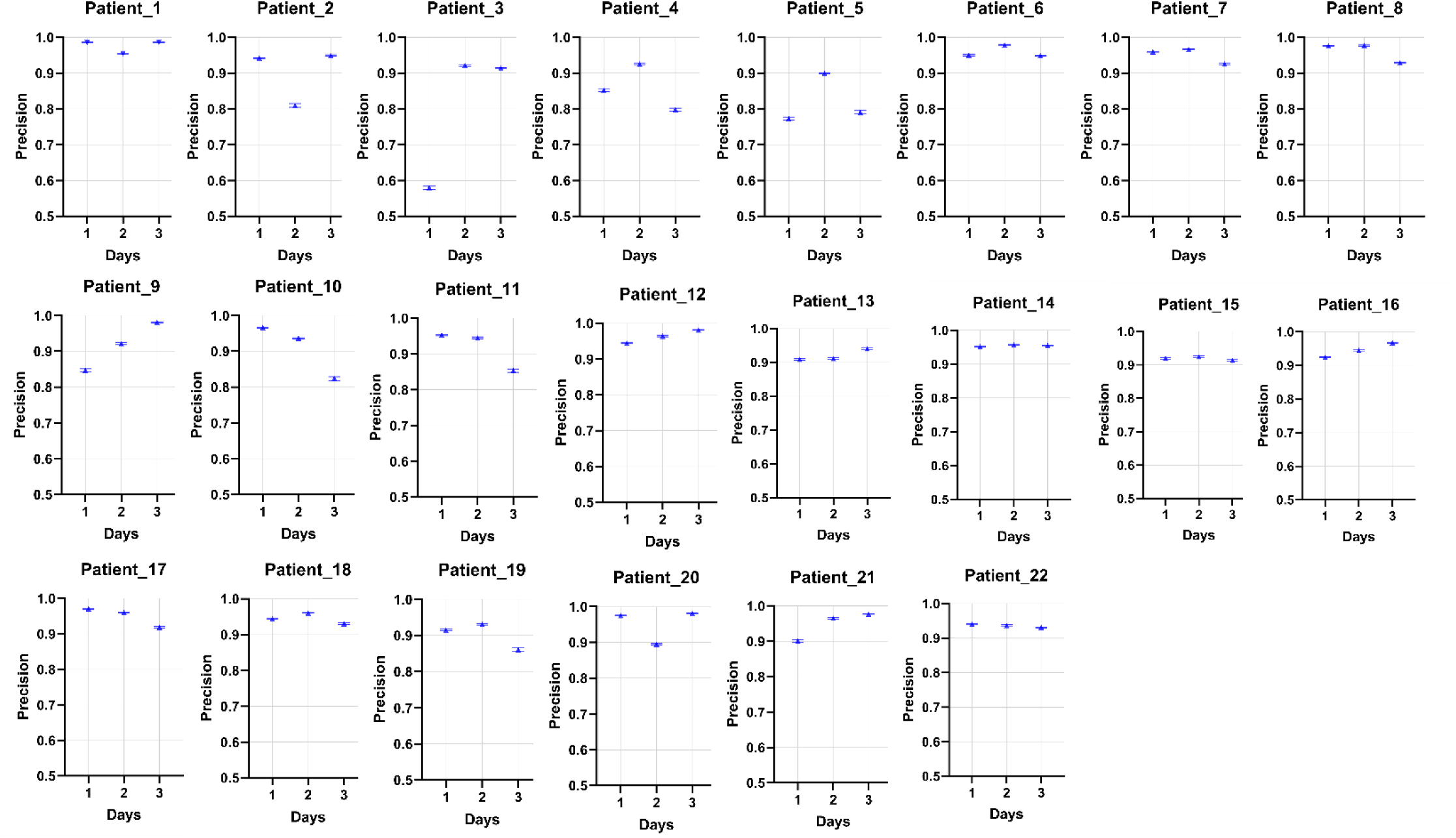
Public Human Precision Values for using Public Human Model. Precision values were evaluated for the low-resolution Public Human pupil dataset using the Public Human Model. For each patient (n = 22) across 3 experimental days, precision and standard error were plotted by day relative to the NeuroPupil model.

**Supplementary Figure 16.**
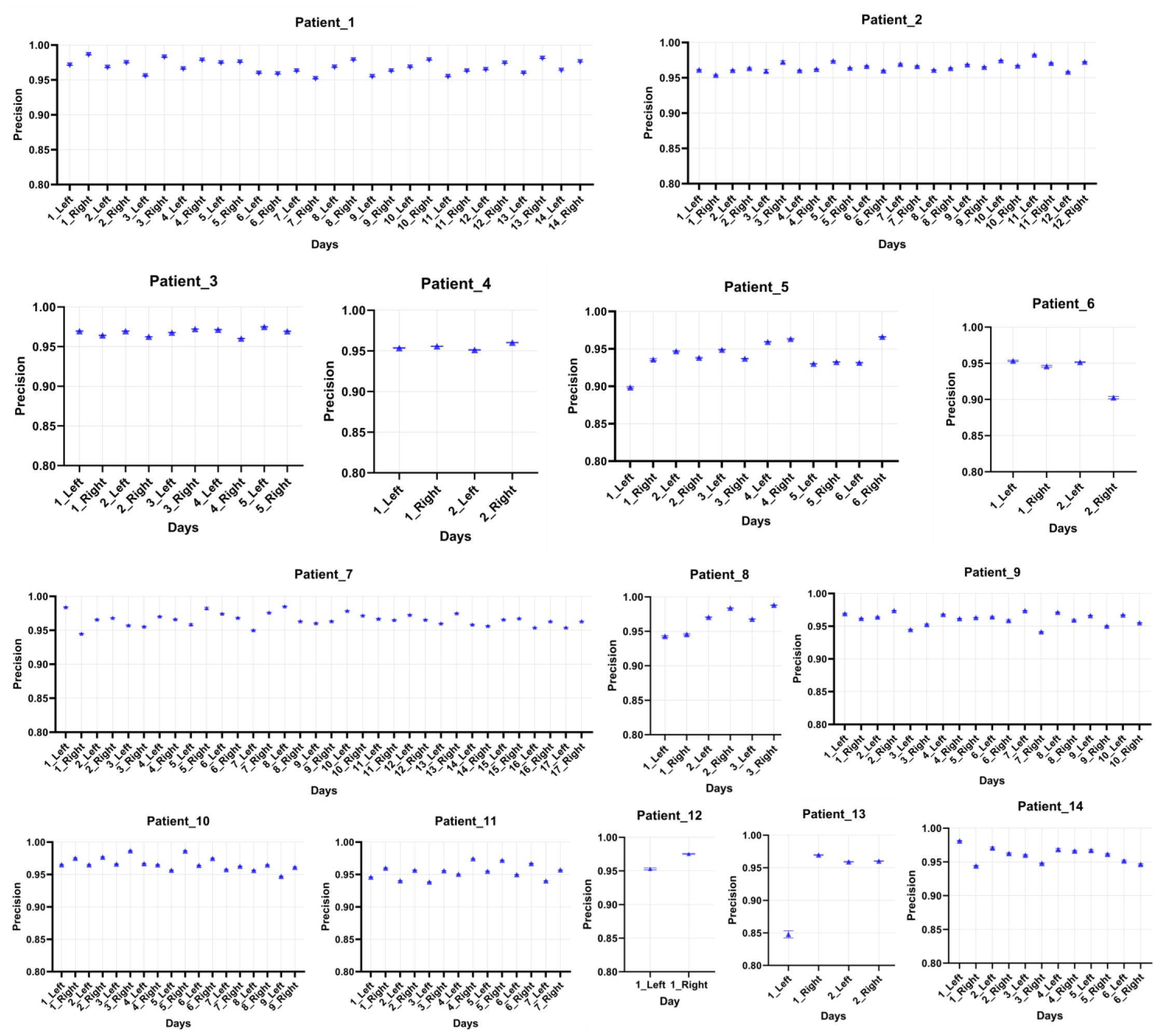
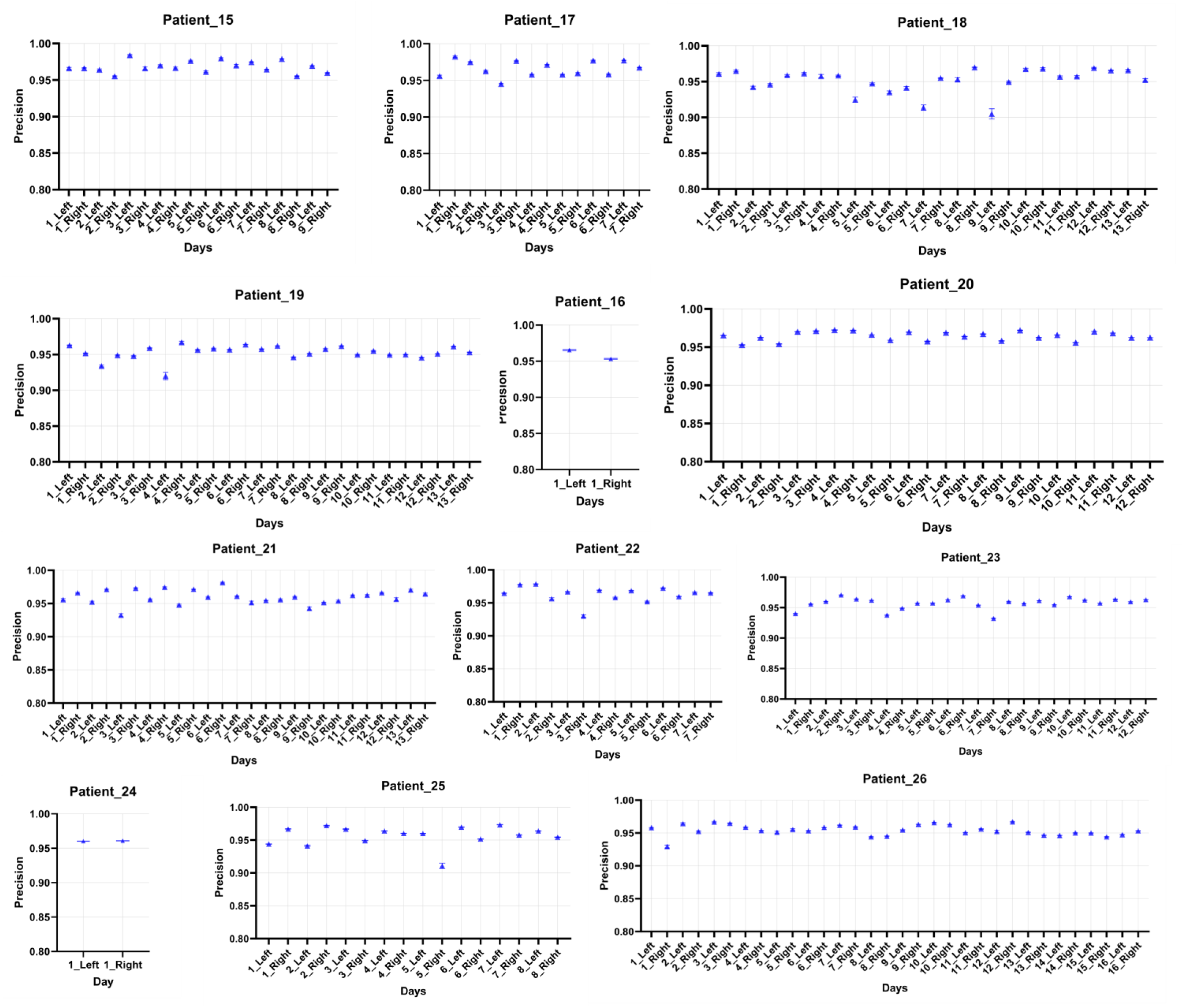
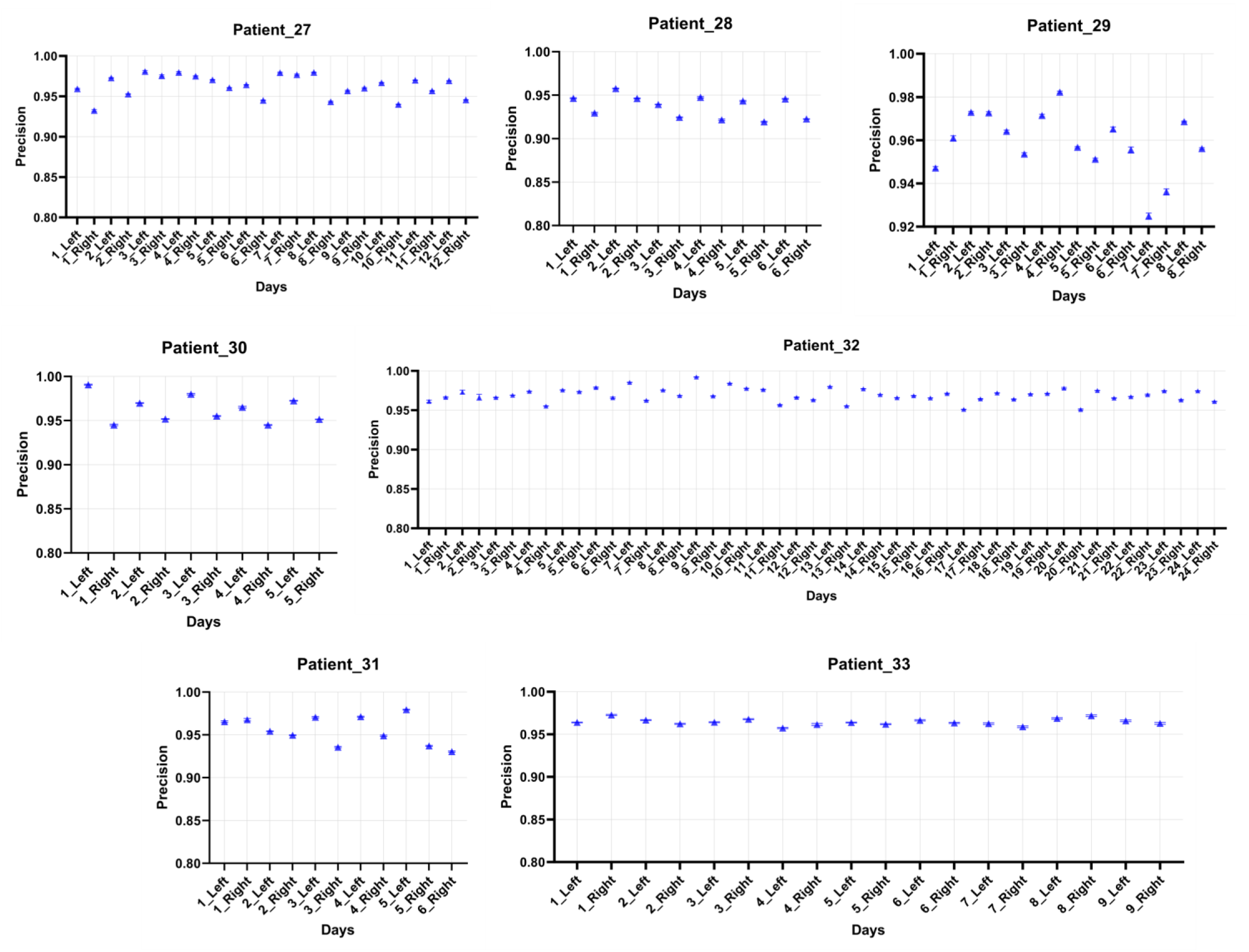
CCF Human Precision Values for using CCF Human Model. Precision values were evaluated for the high-resolution CCF Human pupil dataset using the CCF Human Model. For each patient (n = 33) across a range of experimental days and eye location (Left or Right), precision and standard error were plotted by day relative to the NeuroPupil model.

**Supplementary Figure 17.**
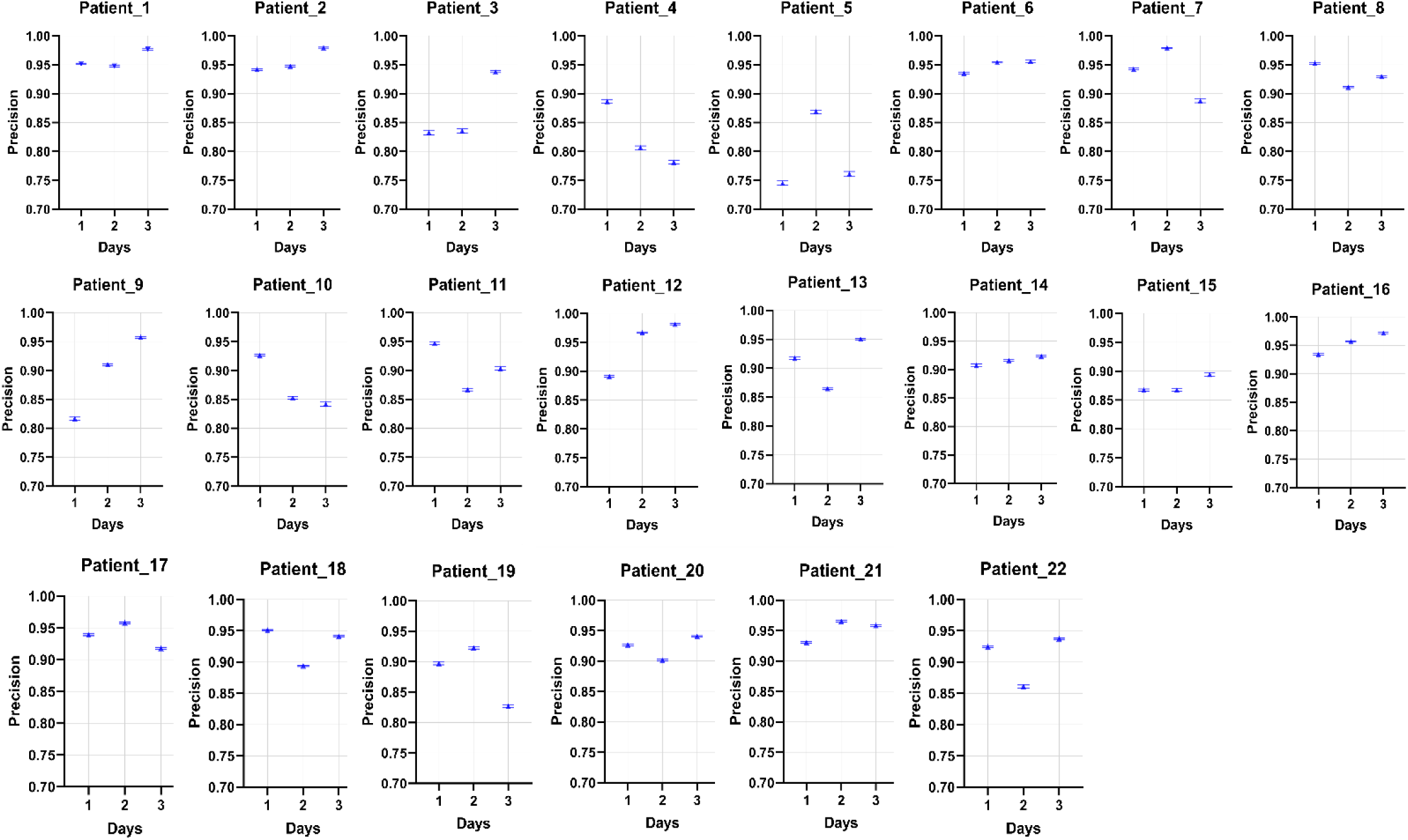
Public Human Precision Values for using NeuroPupil Human Model. Precision values were evaluated for a publicly available low-resolution human pupil dataset using the NeuroPupil Human Model. Each panel corresponds to a single patient (n = 22 patients total), and each point within a panel represents the precision measured on a given experimental day (3 days per patient). Error bars indicate the standard error of precision estimates for that day. Panels are shown separately to illustrate inter-individual variability and within-subject consistency across days.

**Supplementary Figure 18.**
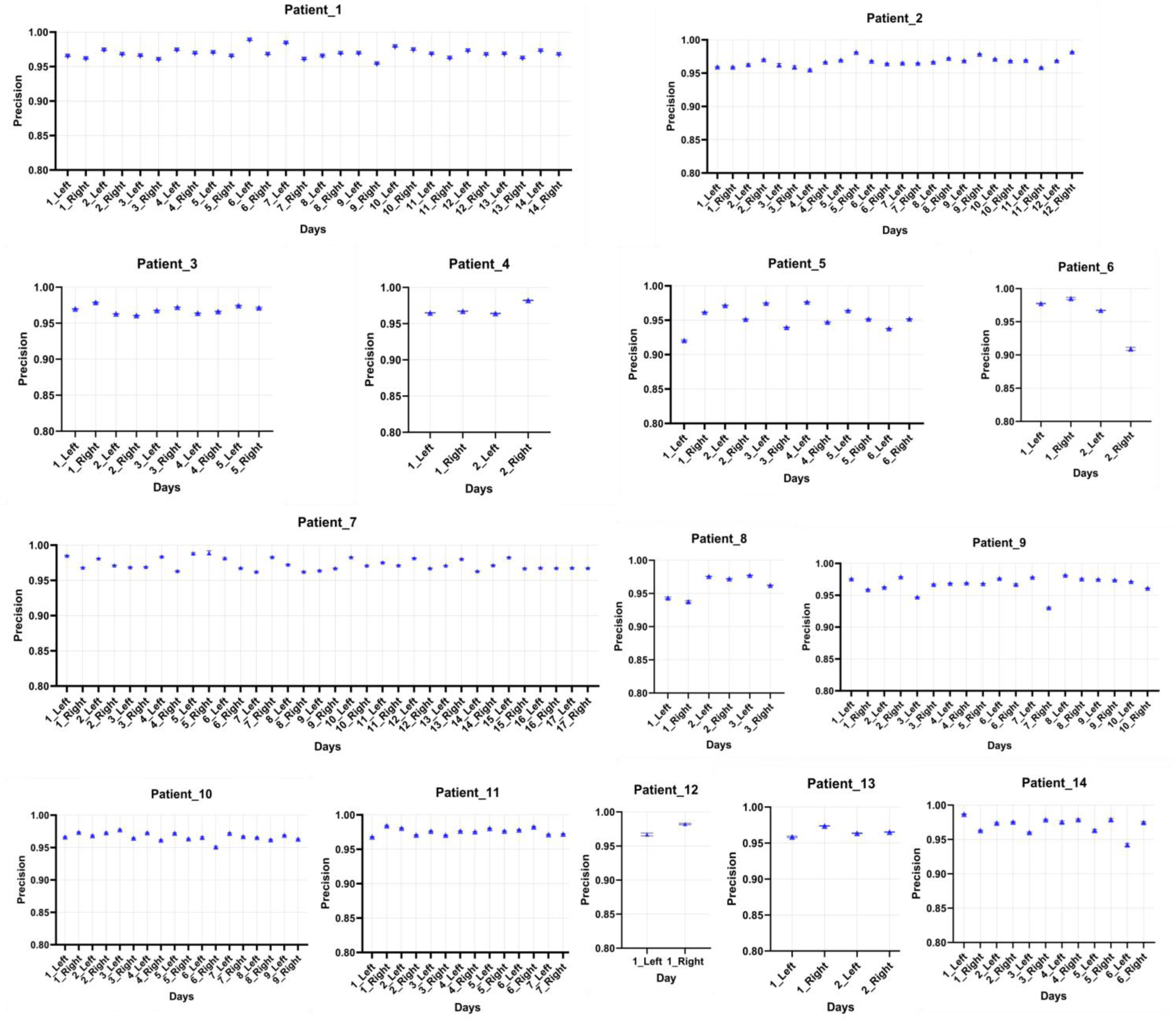
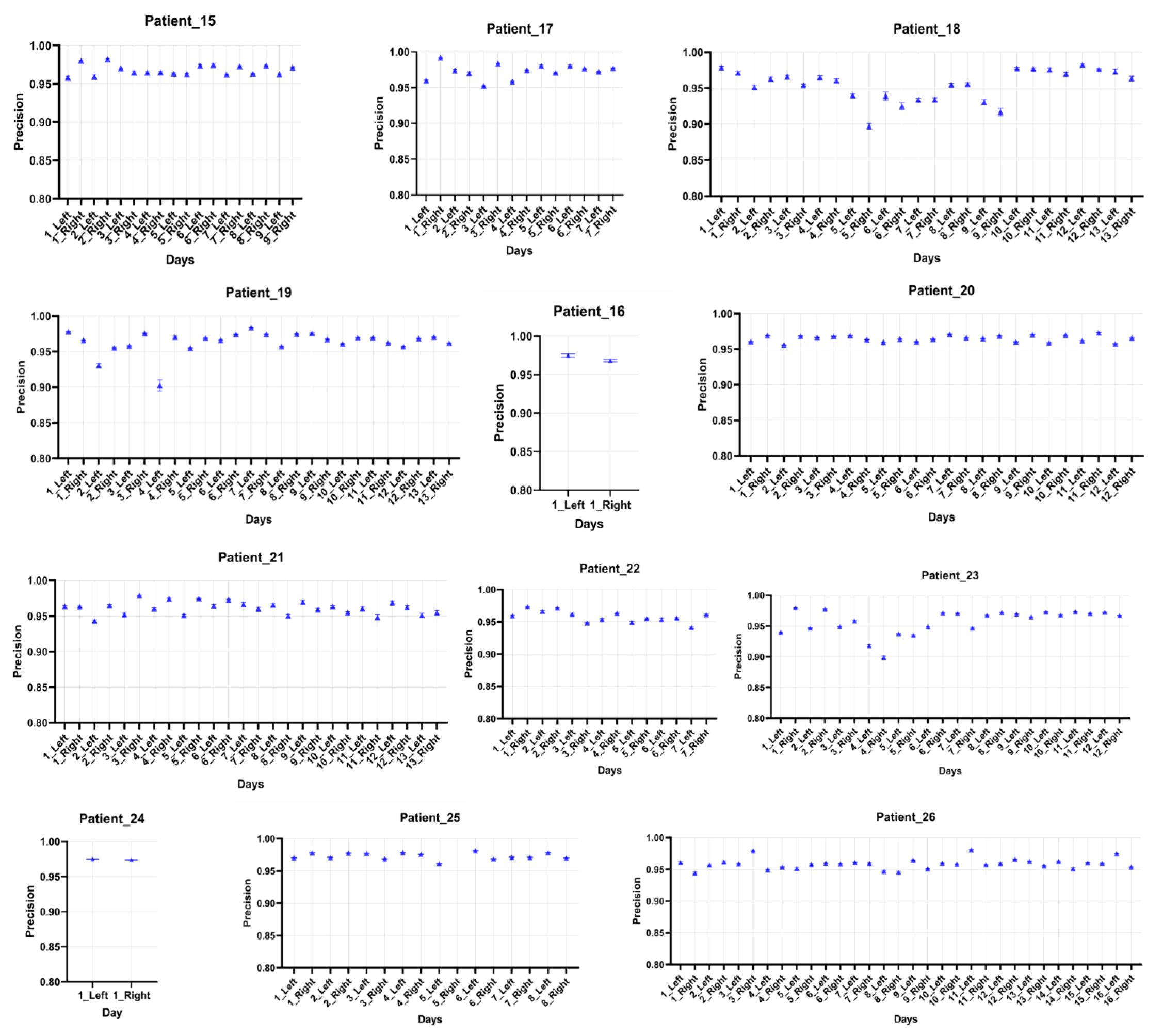
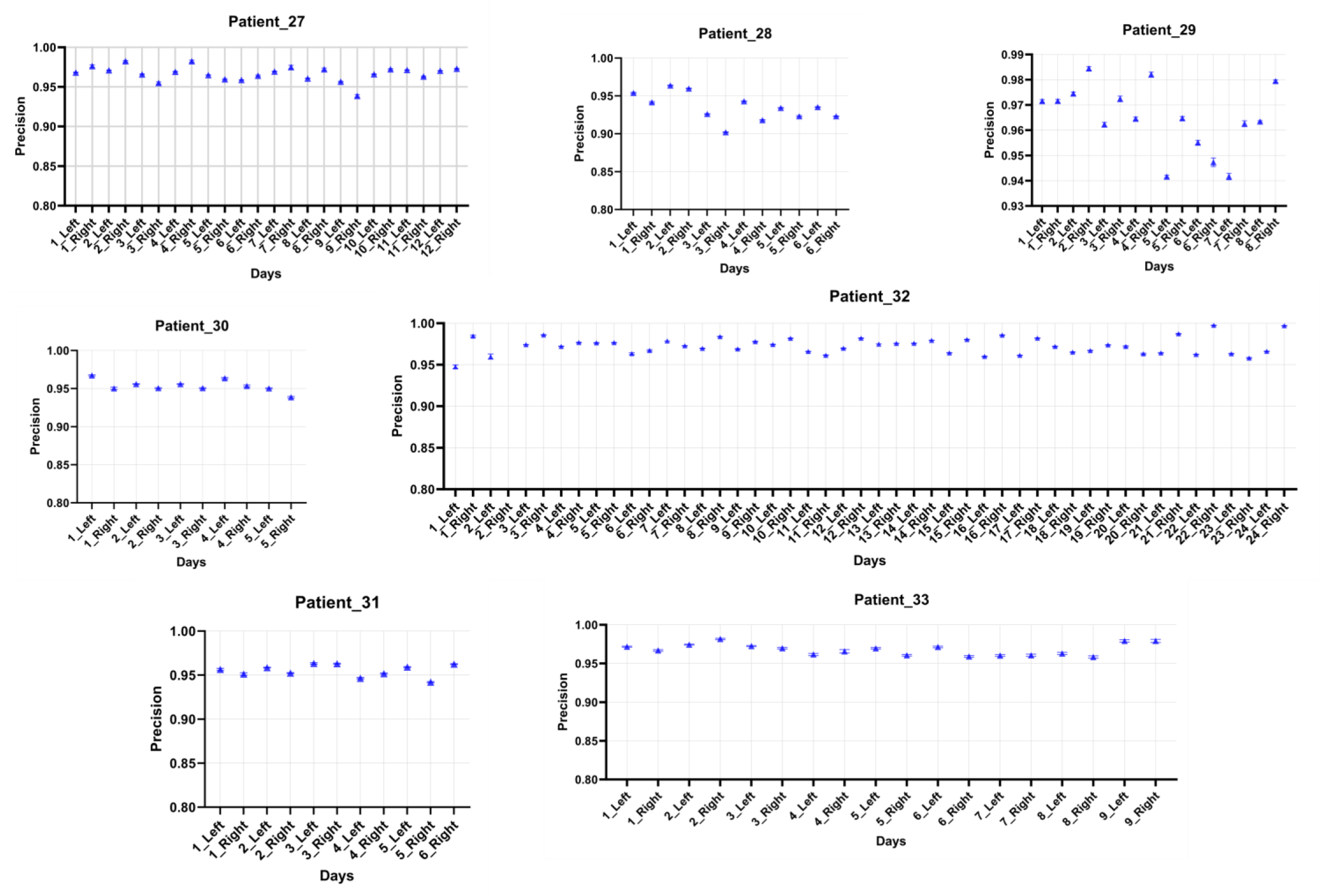
CCF Human Precision Values for using NeuroPupil Human Model. Precision values were evaluated for the high-resolution CCF Human pupil dataset using the NeuroPupil Human Model. For each patient (n = 33) across a range of experimental days and eye location (Left or Right), precision and standard error were plotted by day relative to the NeuroPupil model.

**Supplementary Figure 19.**
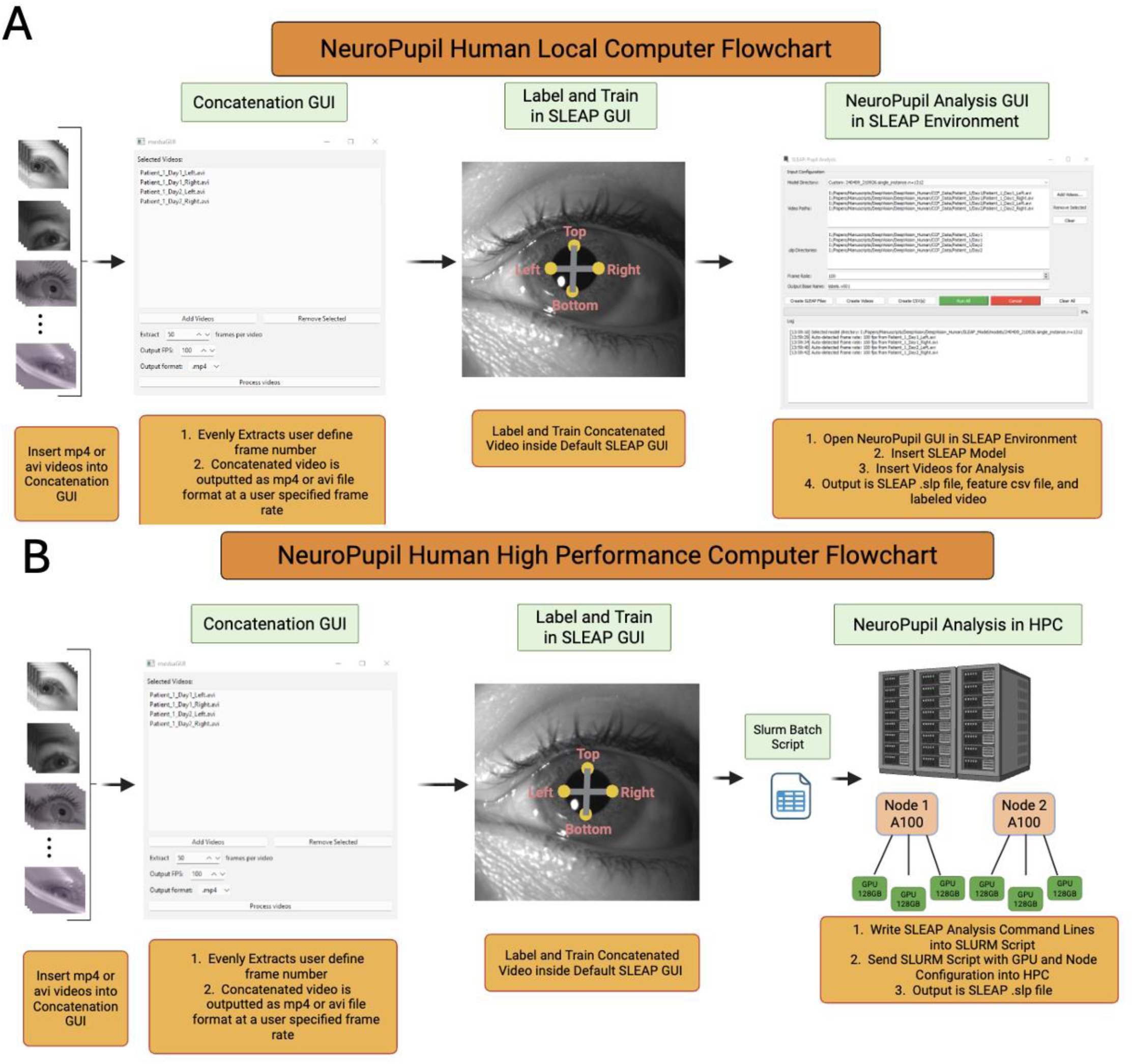
NeuroPupil Workflow for Local and High-Performance Computing Environments Caption. **(A) NeuroPupil Detailed Analysis Flow Chart for Local Computer**. All of the experimental pupil videos (either mp4 or avi video format) are inserted into the Concatenation GUI for creating a concatenated video (mp4 or avi format). The GUI will ask the user to specify the number of frames it will be extracted evenly per video, as well as the concatenated video frame rate and output location. After the video is concatenated, it is uploaded into the SLEAP default GUI for labeling and training the model. After training the model, the custom NeuroPupil Analysis GUI is opened inside the SLEAP Anaconda environment for analyzing the videos. Once the GUI opens, it requires the user to first input the SLEAP model location, then insert the desired videos into the video path location. After the videos are placed into the video path location, the GUI automatically generates the SLEAP analysis file (.slp), feature coordinates csv file, and labeled video inside the same file path as the inserted raw video. Finally, clicking the Run All green button will analyze all of the inserted video data**. (B) NeuroPupil Detailed Analysis Flow Chart for High Performance Computer**. The NeuroPupil analysis process is the exact same as in the local computer until after training the SLEAP model. Once training the model, a slurm batch script is created to analyze the video data. Inside the script, the number of nodes and GPUs are specified, as well as the SLEAP commands to analyze all of the videos. Then, the slurm batch script is sent to the HPC for video analysis and the output is the SLEAP analysis file.

**Supplementary Figure 20.**
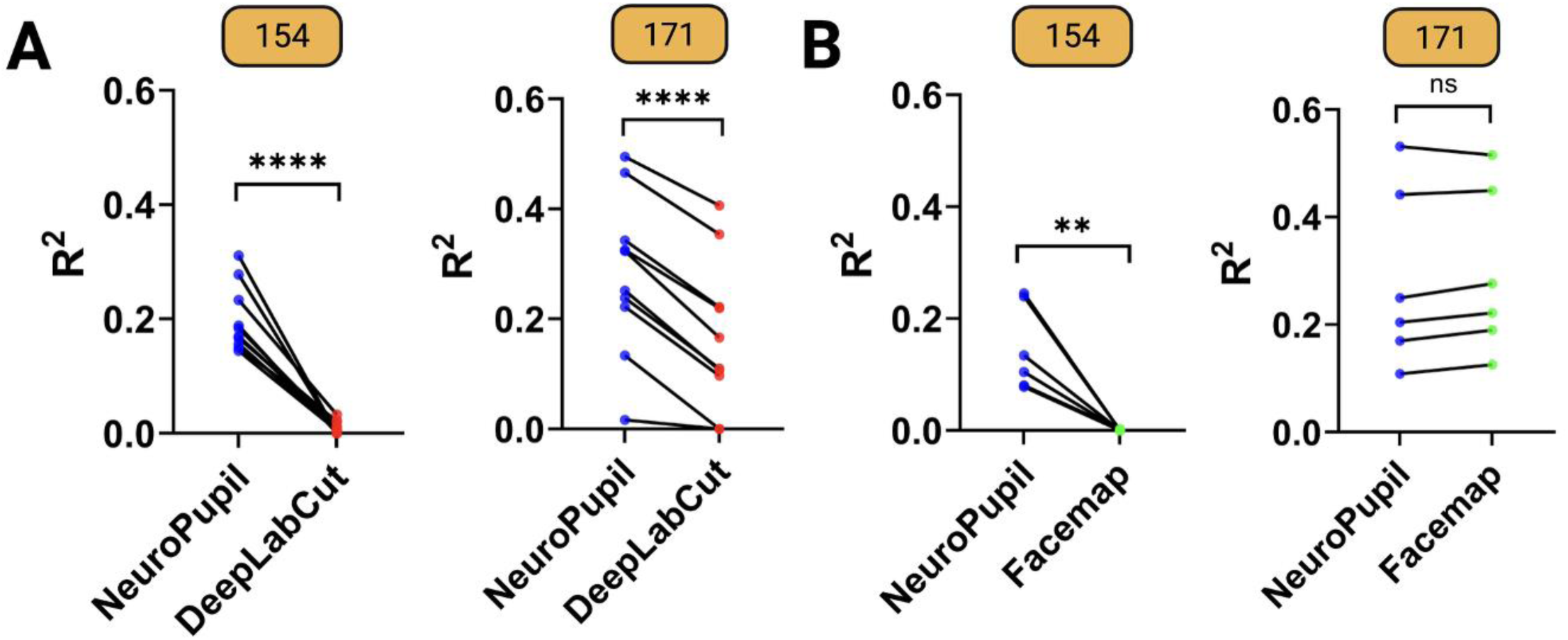
Comparison of Brain Activity Prediction Accuracy (R²) Using Pupil Features. **(A)** Comparison of NeuroPupil and DeepLabCut–based prediction of cortical activity across anatomically defined brain regions. Each panel corresponds to a single animal (Animal IDs 154 and 171 shown; n = 3 animals total). Each dot represents the cross-validated R² value for one brain region, and black lines connect matched regions between methods within the same animal. Predictions were performed using an 8-feature model derived from x/y coordinates of pupil tracking points. Results for Animal 152 are shown in the main figure (Fig. 5B). **(B)** Comparison of NeuroPupil and Facemap–based prediction. Because Facemap outputs only pupil area, a 1-feature prediction model was used. For NeuroPupil, pupil area was computed by ellipse fitting to enable direct comparison. Predictions were therefore limited to six brain regions for which area-based models were evaluated. Each dot represents one region, with lines indicating paired regional comparisons within each animal.

